# The S2 subunit of spike encodes diverse targets for functional antibody responses to SARS-CoV-2

**DOI:** 10.1101/2024.02.26.582219

**Authors:** Jamie Guenthoer, Meghan E. Garrett, Michelle Lilly, Delphine M. Depierreux, Felicitas Ruiz, Margaret Chi, Caitlin I. Stoddard, Vrasha Chohan, Kevin Sung, Duncan Ralph, Helen Y. Chu, Frederick A. Matsen, Julie Overbaugh

**Affiliations:** Human Biology Division, Fred Hutchinson Cancer Center, Seattle, WA 98109, USA; Public Health Sciences Division, Fred Hutchinson Cancer Center, Seattle, WA 98109, USA; Division of Allergy and Infectious Diseases, University of Washington, Seattle, WA 98195, USA; Howard Hughes Medical Institute, Seattle, WA 98195, USA

## Abstract

The SARS-CoV-2 virus responsible for the COVID-19 global pandemic has exhibited a striking capacity for viral evolution that drives continued evasion from vaccine and infection-induced immune responses. Mutations in the receptor binding domain of the S1 subunit of the spike glycoprotein have led to considerable escape from antibody responses, reducing the efficacy of vaccines and monoclonal antibody (mAb) therapies. Therefore, there is a need to interrogate more constrained regions of Spike, such as the S2 subdomain. Here, we describe a collection of S2 mAbs from two SARS-CoV-2 convalescent individuals that target multiple regions in the S2 subdomain and can be grouped into at least five epitope classes. Most did not neutralize SARS-CoV-2 with the exception of C20.119, which bound to a highly conserved epitope in the fusion peptide and showed broad binding and neutralization activity across SARS-CoV-2, SARS-CoV-1, and closely related zoonotic sarbecoviruses. Several of the S2 mAbs tested mediated antibody-dependent cellular cytotoxicity (ADCC) at levels similar to the S1 mAb S309 that was previously authorized for treatment of SARS-CoV-2 infections. Three of the mAbs with ADCC function also bound to spike trimers from HCoVs, such as MERS-CoV and HCoV-HKU1. Our findings suggest there are diverse epitopes in S2, including functional S2 mAbs with HCoV and sarbecovirus breadth that likely target functionally constrained regions of spike. These mAbs could be developed for potential future pandemics, while also providing insight into ideal epitopes for eliciting a broad HCoV response.

**AUTHOR SUMMARY:** The early successes of vaccines and antibody therapies against SARS-CoV-2, the virus responsible for the COVID-19 global pandemic, leveraged the considerable antibody response to the viral entry protein, spike, after vaccination or infection. These initial interventions were highly effective at protecting from infection and reducing severe disease or death. However, SARS-CoV-2 has shown no sign of abating, with the continued rise of new variants that have escaped some of the antibody defense due to distinct alterations most significantly in regions of the spike protein that elicit most of the anti-viral, functional antibody response. These findings suggest a critical need to identify vaccine approaches and therapies that provide the broadest possible antibody responses, focused on regions of spike critical for SARS-CoV-2 infection and, therefore, do not undergo changes that could lead to immune evasion. Our study describes a panel of functional antibodies, from individuals after SARS-CoV-2 infection, that recognize the S2 spike subdomain that is responsible for carrying out viral fusion with host cells. These regions in S2 are generally well conserved across SARS-CoV-2 variants and other closely related viruses and thus, could guide more effective vaccine design in the face of continued viral evolution.

## INTRODUCTION

Eliciting a durable antibody response to SARS-CoV-2 remains an essential component of protection from infection and severe disease from this rapidly evolving virus. So much of the focus on the antibody response has been on antibodies that target the receptor binding domain (RBD) of the spike glycoprotein (1–3), which is a subdomain of S1 and contains the residues that make direct contact with the host cell receptor angiotensin-converting enzyme 2 (ACE2) for viral entry. These mAbs make up the majority of the neutralizing antibody response in vaccinated or convalescent individuals (1, 4–10), making them attractive candidates for further study both as monoclonal antibodies (mAbs) for therapeutics and vaccine design. However, selective pressure on the epitopes of these mAbs has resulted in frequent mutations in RBD of recent VOCs, especially Omicron subvariants (11, 12), dramatically reducing the effectiveness of established responses elicited after vaccination or infection. As a result, vaccine efficacy has been severely compromised against these VOCs (13–16), and the previously approved therapeutic mAbs, which all target the RBD, are no longer effective. Therefore, antibodies that are robust in the face of continued viral evolution and target more conserved regions of spike may be valuable for improving immune durability.

The S2 subunit is a highly conserved region of SARS-CoV-2 spike that has not undergone significant antigenic drift across emerging variants (17). This region, which encompasses the C-terminal half of spike, facilitates fusion of the SARS-CoV-2 envelope with the host cell membrane (18). Several regions of S2 with key roles in this process include the fusion peptide (FP), two heptad repeats 1 and 2 (HR1, HR2) separated by a stem helix linker region (SH), and the transmembrane domain (TM1) (19–22). After the RBD binds to ACE2, spike undergoes a conformational change, which includes a furin cleavage at the S1/S2 boundary, resulting in S1 shedding (3, 23, 24). A second proteolytic cleavage occurs within S2 (S2’ site), primarily via transmembrane serine protease 2 (TMPRSS2) (23, 24), activating a series of conformational changes in S2, exposing the FP and facilitating membrane fusion (25–28). This process is highly conserved across SARS-CoV-2 variants suggesting functional constraints on this process could be keeping the sequences of S2 conserved as well. Studies from other viruses, such as Influenza, Ebola, and human immunodeficiency virus (HIV), have similarly revealed that conserved regions of viral entry proteins involved in membrane fusion are common targets for neutralizing antibodies (29–32), highlighting the need to further explore the role of spike S2 mAbs in protection against SARS-CoV-2.

Despite studies showing that S2-targeting mAbs are an important component of the antibody response in SARS-CoV-2 convalescent patients (33–39), these mAbs remain understudied as compared to RBD mAbs and the range of S2 epitopes they target has not been fully defined. S2 mAbs can exhibit broad neutralization capabilities, and while the potency of these mAbs is lower than their RBD counterparts, their breadth across SARS-CoV-2 VOCs, including Omicron variants, gives them a distinct advantage [reviewed in (40, 41)]. Most neutralizing S2 mAbs target FP (28, 42–44) or SH (44–48) and neutralize predominantly through perturbing membrane fusion as opposed to directly blocking binding of spike to ACE2 (28, 42, 44, 48–50). They have been shown to be protective in animal models of SARS-CoV-2 infection as well (42, 50, 51). S2-targeting mAbs make up a considerable amount of the binding IgGs in SARS-CoV-2 vaccinated and convalescent individuals (1, 52), yet as mentioned, they comprise only a small percentage of the neutralizing IgGs; therefore, S2 antibodies could be playing a non-neutralizing role as well. Recent studies increasingly suggest a role for protective non-neutralizing antibodies in animal models where neutralization and protection are not correlated (53, 54). In these models, ablation of Fc receptor (FcR) binding of experimental mAbs led to reduced efficacy, supporting a role for FcR-mediated function in antibody protection against SARS-CoV-2 (53). Even neutralizing mAbs, such as S309 that was previously authorized for treatment of SARS-CoV-2 (49, 55), have been shown to rely on intact Fc-effector function to achieve optimal in vivo protection (56). Several S2 mAbs that mediate Fc-effector functions, such as antibody-dependent cellular cytotoxicity (ADCC) or cellular phagocytosis (ADCP) have been described (45, 50, 57, 58). The mAb S2P6, for example, can promote ADCC in an *in vitro* assay with levels on par with S309 (55). The Fc-effector function for S2 mAbs is also an essential component of their activity *in vivo* in animal systems including protection from infection and reducing severe disease post-infection (45, 50, 57, 58).

The highly conserved S2 region is also a preferential target for antibodies that demonstrate broad cross-reactivity against related human coronaviruses (HCoVs) (35, 37, 44, 59, 60). SARS-CoV-1 and SARS-CoV-2 both belong to the sarbecovirus family, which is comprised of a collection of viruses residing in animal reservoirs capable of binding to human ACE2 with the potential for spillover (61). In addition, SARS-CoV-1 and SARS-CoV-2 are two of seven HCoVs known to cause human disease. Four HCoVs are responsible for mild infections associated with the common cold including alphacoronaviruses HCoV-229E and HCoV-NL63, and betacoronaviruses HCoV-OC43 and HCoV-HKU1; and three betacoronaviruses that cause more severe disease in humans (MERS-CoV, SARS-CoV-1, and SARS-CoV-2). The S2 subunit, especially the FP region, has upwards of 63-98% sequence similarity between HCoVs (62). Studies from early in the SARS-CoV-2 pandemic identified cross-reactive antibody responses to the S2 region in uninfected individuals prior to the development of the vaccine suggesting there was expansion of memory B cells that were initially elicited against endemic HCoVs with cross-reactivity to SARS-CoV-2 (36, 39, 63–65). Further, several of the reported S2 mAbs isolated from SARS-CoV-2 convalescent individuals have demonstrated broad binding to HCoVs and sarbecoviruses and in some cases broad neutralizing activity as well (35, 66).

In this study, we describe a collection of S2 mAbs isolated from two individuals following SARS-CoV-2 infections. A subset of these mAbs demonstrated antiviral activity, such as neutralization and Fc-effector function in the form of ADCC. Further, some of these mAbs exhibited broad reactivity to HCoVs (both beta- and alphacoronaviruses) and closely related sarbecoviruses. These mAbs targeted diverse regions of S2, some that presented as linear peptides including the highly conserved and immunogenic FP region, and others that likely bind less well described conformational epitopes.

## RESULTS

### Isolation of mAbs that target non-S1 regions of the spike glycoprotein in SARS-CoV-2 convalescent individuals

To identify mAbs that target more conserved regions of spike outside of the S1 subunit in the S2 region, we isolated spike-specific memory B-cells from PBMC samples collected thirty days post-symptom onset (dpso) from two individuals recovering from SARS-CoV-2 infections. Antibody responses to spike and the S2 subunit tend to be more robust in individuals who have experienced a SARS-CoV-2 infection compared to vaccination alone, and particularly in those with hybrid immunity from combined vaccination and infection (38, 67), and we focused on individuals with these types of exposure histories to isolate S2 mAbs. One participant, C20, was infected with the original Wuhan-Hu-1 (WH-1) strain prior to the availability of SARS-CoV-2 vaccines. The other participant, C68, had been vaccinated with two doses of the Pfizer-BioNTech mRNA vaccine and experienced a Delta VOC breakthrough infection two months after receiving the second dose. Memory B cells were isolated from PBMCs by single cell sorting with a bait approach to enrich for spike-specific B cells. From the C20 PBMCs, we collected memory B cells that were enriched for binding to WH-1 S2 protein; and for C68, memory B cells were sorted based on binding to WH-1 S2 protein and/or Delta prefusion-stabilized spike trimer, which were both used as bait (68).

mAbs with paired heavy (IgH) and light (IgL) chain sequences that had in-frame variable regions were generated and assayed for binding to prefusion-stabilized WH-1 spike trimer and WH-1 S1 and S2 monomers to determine their specificity. For this initial binding screen, we tested all mAbs in an ELISA at a single concentration (1 μg/mL) that was saturating for most antibodies that bind WH-1 Spike trimer (**Fig 1**). We included mAbs with known binding specificity for S1 (LY-CoV-1404 (69)) or S2 (mAb B6 (70)), and a non-spike, HIV mAb VRC01 (71) as controls. We isolated 11 mAbs from C20 that bound to spike trimer and/or the S2 monomer, and 100 mAbs from C68 that bound to at least one of the three antigens (spike trimer, S1, or S2). For this study, to identify mAbs with S2 epitopes we focused on mAbs that did not bind to the S1 protein (OD450nm < 0.18, threshold defined OD450nm_ave_ of the HIV mAb VRC01 + 3 standard deviations, **S1 Fig**), and across C68 and C20 mAbs we identified 40 that are likely S2-specific (**Fig 1**).

**Fig 1.**
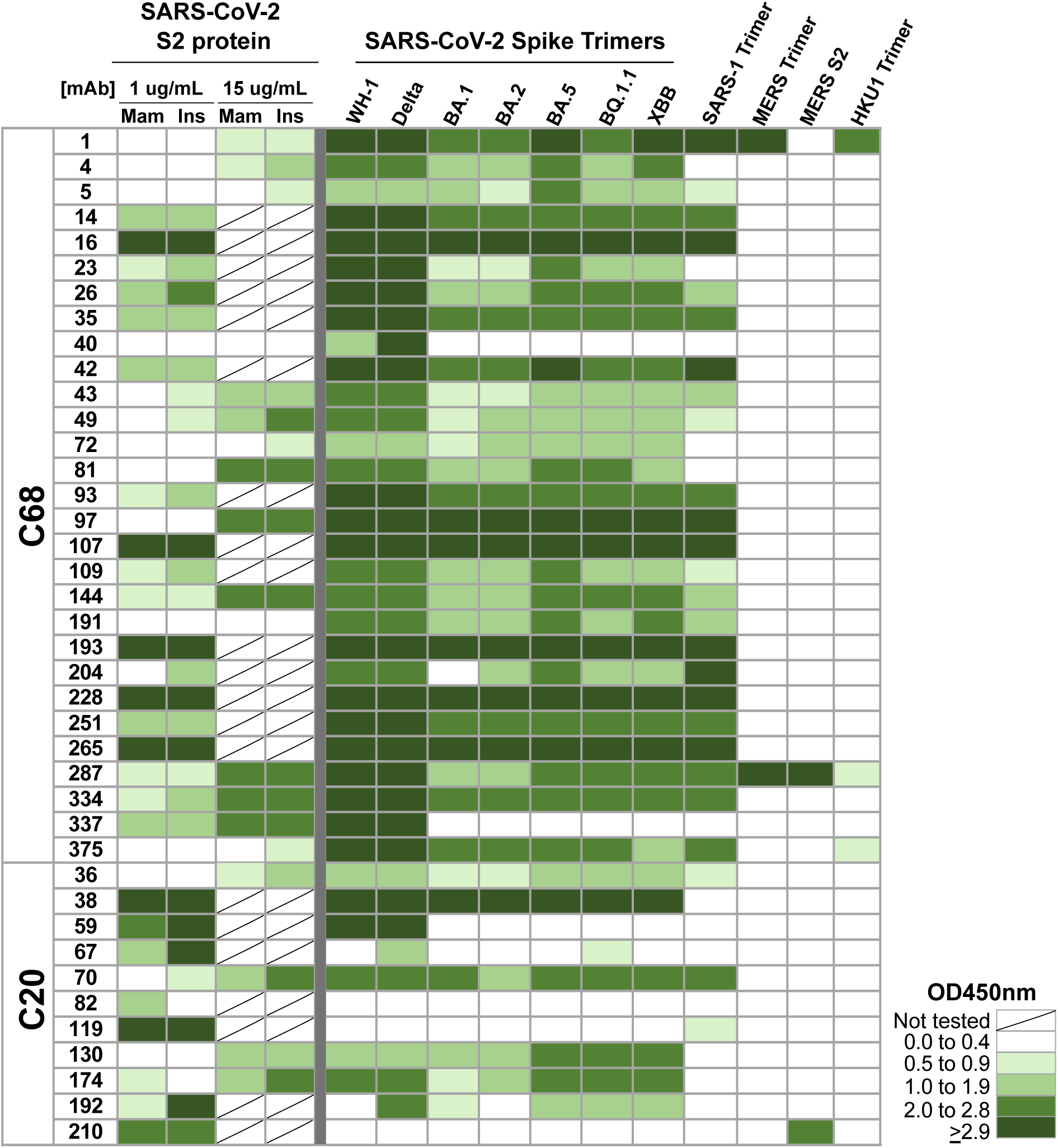
Binding profiles of S2-targeting mAbs from SARS-CoV-2 convalescent cases. Heatmap showing binding of mAbs from C68 and C20 (rows) to spike antigens by ELISAs. Spaces are colored based on the OD450nm values with higher numbers and more binding represented in darker green. Antigens (columns) included the following: SARS-CoV-2 WH-1 S2 subunit protein produced in mammalian cells (Mam) or insect cells (Ins), SARS-CoV-2 pre-fusion stabilized spike trimers from WH-1 and VOCs (Delta, Omicron VOCs BA.1, BA.2 BA.5, BQ.1.1, XBB), SARS-CoV-1, MERS-CoV, HCoV-HKU1, and MERS-CoV S2 monomer protein. All mAbs were tested at 1 μg/mL, except against the SARS-CoV-2 S2 protein the mAbs were tested at 1 μg/mL or 15 μg/mL as noted. Values were averaged over two technical replicates.

We evaluated binding to the S2 subunit in multiple ways because many of the mAbs bound S2 protein weakly in the initial screen (OD450nm ≤ 1.0) making it difficult to definitively verify their specificity (**Fig 1, S1 Fig**). First, we tested all 40 mAbs for binding to S2 protein produced in baculovirus-infected insect cells (72) as compared to the same protein produced in mammalian HEK293 cells used in the initial screen. Insect cell-expressed proteins can have less complex or otherwise altered glycan structures compared to mammalian-expressed proteins (73). Therefore, some S2 epitopes could be more accessible in the insect-produced S2 in this ELISA assay. Almost all of the mAbs demonstrated similar or increased binding to the insect-produced S2 compared to the mammalian-produced S2 protein (**Fig 1, S1 Fig**). We further interrogated 19 mAbs with weak binding to both S2 proteins (OD450nm < 1 for both antigens) using a higher concentration of antibody (15 μg/mL). Even at the higher concentration, none of the 19 mAbs bound to S1 protein above background (OD450nm_ave_ of the HIV mAb VRC01 + 3 standard deviations) (**S1B Fig**). Almost all of the mAbs exhibited increased binding to an S2 protein at the higher concentration compared to 1 μg/mL (**Fig 1**). Six of the mAbs demonstrated detectable, but weak binding to S2 proteins (OD450nm > 1), and one mAb, C68.40, did not bind S1 or S2 proteins (**Fig 1, S1B Fig**). All of these mAbs bound spike trimer well suggesting their epitopes may depend on the trimer conformation and thus would not bind well to unstable S2 monomer alone. However, since they did not bind S1, we have included these mAbs in this set of likely S2 mAbs for further characterization.

These 40 mAbs were also tested for binding against a panel of spike trimers representing different SARS-CoV-2 VOCs and HCoVs at 1 μg/mL (**Fig 1**). Many of the C68 S2 mAbs and some of the C20 mAbs demonstrated binding breadth across the SARS-CoV-2 VOCs including XBB and BQ.1.1. Further, most of those broad mAbs also bound SARS-CoV-1 spike trimer. Four of the mAbs bound to spike antigens from other betacoronaviruses: C68.1 bound to MERS-CoV and HCoV-HKU1 spike trimers, C68.287 bound near saturation to MERS-CoV spike trimer and S2 protein and weakly to HCoV-HKU1 trimer, C68.375 bound weakly to HCoV-HKU1 spike, and C20.210, bound to the S2 subunit of the MERS-CoV spike suggesting these mAbs target a highly conserved region of S2.

### V(D)J Gene Usage and features

We next evaluated the V(D)J gene usage across the 40 S2 mAbs described above. To identify the inferred germline genes and annotate gene features, we used *partis*, a software package specifically designed for this purpose (74–76). We noted that a few germline genes were shared between individuals C20 and C68, such as the IGHV4 genes, but many of the genes were differentially used between individuals (**Fig 2A-C, S1 Table**). For the heavy chain gene (IgH), C68 mAbs were enriched for IGHV1-69, IGHV3-30, IGHV3-7 whereas C20 mAbs used IGHV3-30-3 more frequently (**Fig 2A**). On the whole, the gene usage patterns in these mAbs were consistent with clonotypes that have been observed in other studies of SARS-CoV-2 spike-specific mAbs (40, 77–79) especially for S2-targeting mAbs, which tend to have higher usage of IGHV1-69, IGHV3-30, IGHV3-30-3 (50, 79–82). For the light chain genes (IgL) encompassing IGK and IGL genes, we again observed enriched usage of genes (e.g., IGLV3-10, IGKV3-11, IGK3-20) that have previously been observed in other SARS-CoV-2 mAbs (50, 81) (**Fig 2B-C**).

**Fig 2.**
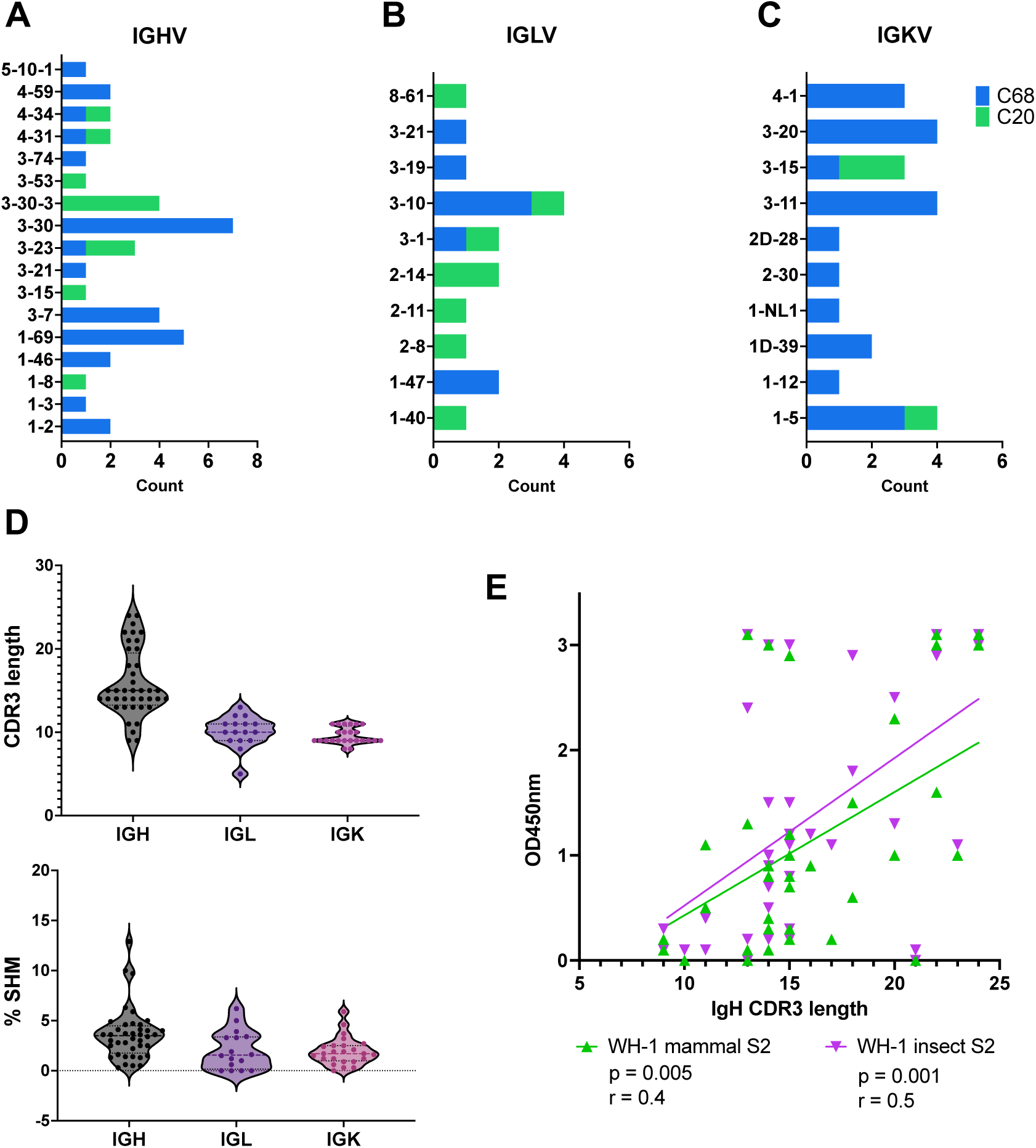
Heavy and light chain V(D)J gene usage and features of mature S2 mAbs. Diversity of the germline V gene usage for *IGH* **(A)**, *IGL* **(B)**, and *IGK* **(C)** for C68 (blue) and C20 (green) S2 mAbs. (**D**) Distribution of CDR3 amino acid (aa) length (top) and % SHM (bottom) in *IGH*, *IGL*, and *IGK* genes. **(E)** Correlation plot comparing CDR3 aa length in *IGH* to binding of mAb to WH-1 S2 protein expressed in mammalian cells (green) or insect cells (purple). Binding was determined by ELISA and shown as background-corrected OD450nm values. Pearson’s correlation analysis was used to determine the strength of the correlations.

We also assessed two mAb gene features, percent somatic hypermutation (%SHM) and length of the complementarity-determining region 3 (CDR3), calculated with *partis*. The CDR3 length ranged from 9-24 amino acids and 5-13 amino acids for the IgH and IgL genes, respectively (**Fig 2D, top**). The majority of the mAbs had relatively low %SHM (<5%) in the IgH and IgL genes (**Fig 2D, bottom**). We looked for associations between CDR3 length or %SHM and binding to antigen and observed positive correlations between the IgH CDR3 length and binding to the WH-1 S2 protein (mammalian S2 p=0.005; insect S2 p=0.001) (**Fig 2E**). The IgL CDR3 length was not correlated with S2 binding (**S2 Fig**). There were no significant correlations observed between CDR3 length and binding to WH-1 spike trimer (**S2 Fig**) and between %SHM and binding to any of the antigens (**S2 Fig**).

### Neutralizing activity of S2 mAbs

To identify S2 mAbs that exhibit neutralization capabilities, we evaluated their potency against spike-pseudotyped lentiviruses that can infect HEK293T-ACE2 cells, used previously (68, 83). All of the S2 mAbs were screened at 10 μg/mL for the ability to neutralize SARS-CoV-2 WH-1, which was the strain both individuals were exposed to, through infection for C20 and the original vaccine for C68. Only C20.119 was able to neutralize WH-1 greater than 50% neutralization and showed 80% virus neutralization (**Fig 3A**). Notably, among the S2 mAbs identified here, C20.119 had the highest IgH %SHM (12.9 %SHM). We then evaluated the ability of C20.119 to neutralize a panel of SARS-CoV-2 VOCs using full mAb dilution curves to calculate IC50s. C20.119 was able to neutralize WH-1 and Delta VOCs with comparable IC50s (1.1 and 2.4 μg/mL, respectively, **Fig 3B**). Against the Omicron VOCs, C20.119 was still able to neutralize but it was ∼10 to 60-fold less potent (**Fig 3B, 3D**).

**Fig 3.**
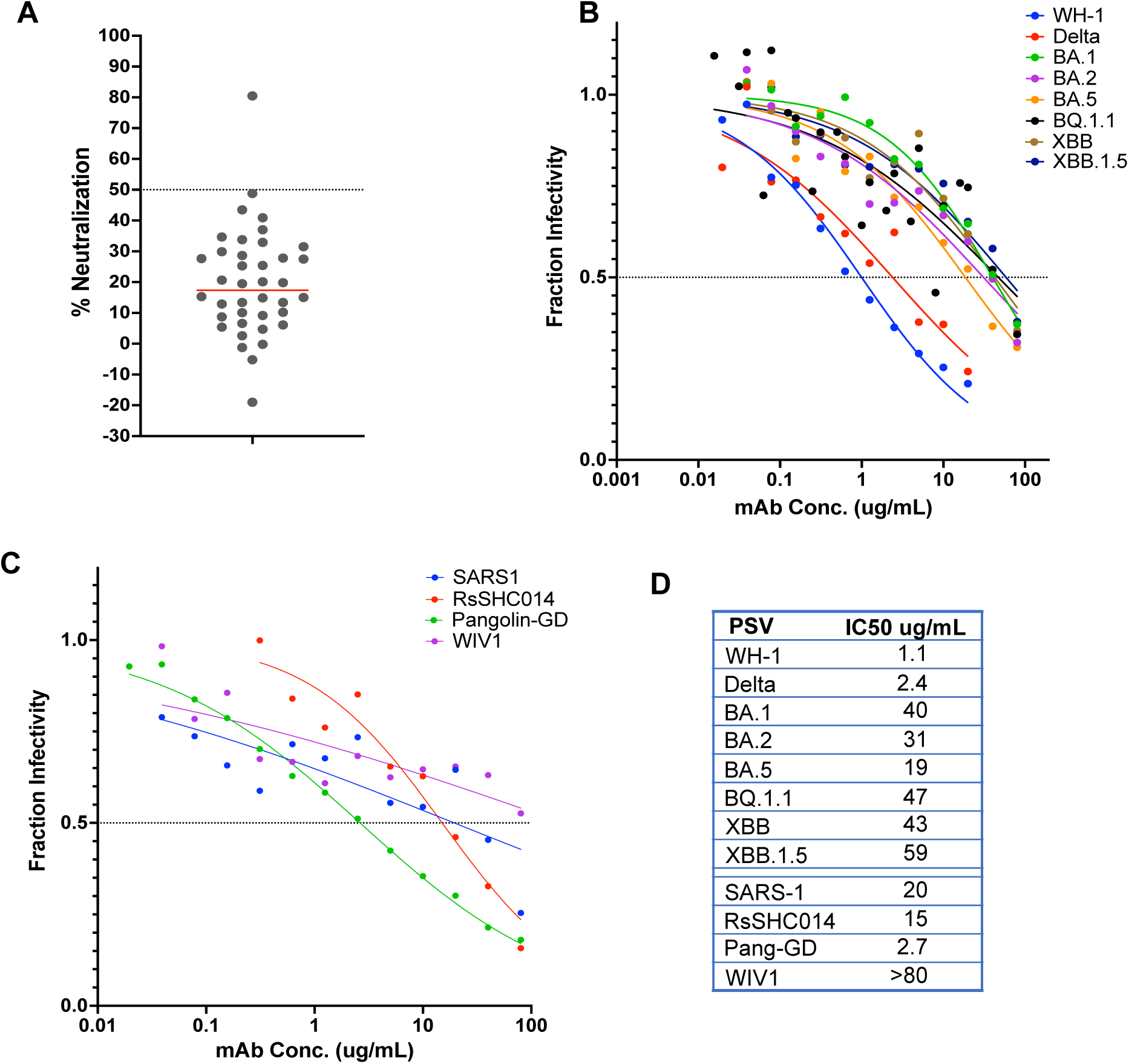
Neutralization potency and breadth of S2 mAb C20.119 against SARS-CoV-2 VOCs, SARS-CoV-1, and related zoonotic sarbecoviruses. **(A)** Screening of all S2 mAbs for neutralization of WH-1 spike-pseudotyped lentiviruses. All antibodies were tested at 10 μg/mL and the target cells were ACE2-expressing HEK293T cells. The percent neutralization of the pseudoviruses is shown, averaged across two technical replicates. One mAb, C20.119, was able to neutralize greater than 50%, which was the threshold for further assessment, shown by the dashed line. Neutralization of a panel of SARS-CoV-2 variant pseudoviruses **(B)** or sarbecovirus pseudoviruses **(C)** by C20.119. SARS-CoV-2 pseudoviruses tested included WH-1, Delta, Omicron VOCs BA.1, BA.2, BA.5, BQ.1.1, XBB, XBB.1.5. The related sarbecovirus pseudoviruses assessed were SARS-CoV-1, bat viruses RsSHC014 and WIV1, and Pangolin-GD. C20.119 was tested in 12-point dilution curves starting at 20 μg/mL for WH-1 and Delta and 80 μg/mL for the other pseudoviruses. Fraction infectivity values are the average of two technical replicates in two to three independent experiments using at least two pseudovirus batches. **(D)** Pseudovirus neutralization IC50 values for C20.119.

We also tested whether C20.119 could neutralize viruses pseudotyped with spikes from SARS-CoV-1 and other closely-related, zoonotic sarbecovirus strains. The SARS-CoV-2 WH-1 and SARS-CoV-1 spike sequences are 86% similar (84), but the S2 subdomain sequences are ∼90% conserved and several reported S2 neutralizing antibodies have broad activity across the two sarbecoviruses (85). C20.119 neutralized SARS-CoV-1 with an IC50 of 20 μg/mL, on par with its potency against the SARS-CoV-2 VOCs (**Fig 3C, 3D)**. We selected three representative sarbecoviruses to test based on their relatedness to SARS-CoV-1 and SARS-CoV-2 and their ability to bind human ACE2 (86). Pangolin-GD found in pangolins of the Guangdong Province, is overall ∼90% similar to SARS-CoV-2 spike (87, 88), and two bat viruses RsSHC014 and WIV1, approximately 93% and 74% similar to SARS-CoV-2 spike, respectively (84, 88, 89). C20.119 was able to neutralize RsSHC014 (IC50 = 20 μg/mL) and Pangolin-GD (IC50 = 5.3 μg/mL) comparable to its potency against SARS-CoV-2. Against WIV1, while there was a dose response, the IC50 was above the maximum concentration of 80 μg/mL. Overall, C20.119 demonstrated neutralization breadth across SARS-CoV-2 VOCs and closely related sarbecovirus, though was only weakly potent.

### S2 mAbs mediate antibody-dependent cellular cytotoxicity (ADCC)

To identify S2 mAbs that might exhibit non-neutralizing functions against SARS-CoV-2, we assessed the ability of these mAbs to mediate ADCC, which has been identified as a correlate of protection for SARS-CoV-2 in both humans and animal models (54, 90, 91). ADCC activity was examined using primary human effector cells (i.e., PBMCs) and target cells which are resistant to direct cell killing by natural killer (NK) cells and stably express the full length, GFP-tagged WH-1+D614G spike glycoprotein (CEM.NKr.Spike) (92). We selected a subset of the S2 mAbs based on their ability to bind WH-1 spike trimer in ELISAs and their binding breadth across SARS-CoV-2 VOCs and SARS-CoV-1 (**Fig 1**). Prior to performing the ADCC experiment, we confirmed that all 17 of the mAbs were able to bind to the Spike protein expressed by the target cells with the same parameters to be used in the ADCC assay. All S2 mAbs could bind to the CEM.NKr.Spike cells (∼100% of them were stained), with variable MFI (**S3 Fig**). As a measure of ADCC, we quantified NK cell (CD56+CD3-) activation (surface CD107a and intracellular IFN-γ) by flow cytometry (gating strategy shown in **S4A Fig)** following incubation of PBMCs from healthy donors (containing NK cells), spike-expressing target cells, and mAbs, a standard method for evaluating ADCC function of SARS-CoV-2 mAbs (93–95). In addition, we measured CEM.NKr.Spike target cell death to further evaluate the non-neutralizing function of the S2 mAbs (gating strategy shown in **S4B Fig**). We included the RBD mAb S309, which was previously authorized for treatment of SARS-CoV-2 infections and has both neutralization and ADCC function *in vitro* and *in vivo* (49, 55). The mean fluorescence intensity (MFI) of the S2 mAbs binding to CEM.NKr.Spike cells did not correlate with their ability to activate NK cells (%CD107a+, p = 0.8) (**S3D Fig**), suggesting S2 mAb binding does not influence ADCC.

With Donor 1 PBMCs, all of the mAbs lead to NK-cell activation above background and the negative control HIV-specific mAb A32 ranging from 1.8 to 35.6% (**Fig 4A**). Five mAbs lead to activation of over 20% of NK cells (C68.16, C68.107, C68.287, C20.36, C20.130), which was on par with the activation seen with previously approved therapeutic mAb S309 (%CD107a+ NK cells = 23.5%). The mAb mediating the highest NK cell activation was C20.36 at 35.6% CD107a+ NK cells. C20.36 also resulted in the highest %IFN-γ+ NK cells (7.8%) and target cell death (7.9%) (**S5 Fig**). Across the set of mAbs, the %IFN-γ+ NK cells and %CEM.NKr.Spike target cell death were highly correlated with the %CD107a+ NK cells (**Fig 4B & 4C**). Across the 17 S2 mAbs tested, the overall NK cell response was more robust with Donor 1 PBMCs versus Donor 2 PBMCs, but the percentages of CD107a+ NK cells with each mAb were highly correlated between the donors (**S6 Fig**, r=0.87, p<0.0001). While C20.36 had the highest ADCC activity, it demonstrated limited binding breadth across the SARS-CoV-2 VOCs (**Fig 1**), as was the case for C20.174 and C20.130 that also had high ADCC within these mAbs. In addition, C68.16, C68.107, C68.1, and C68.287 that all mediated 15-20% NK cell activation exhibited significant binding breadth across SARS-CoV-2 VOCs, SARS-CoV-1, and the latter two bound other HCoVs as well (**Fig 1**) indicating a set of mAbs that could have functional activity against HCoVs.

**Fig 4.**
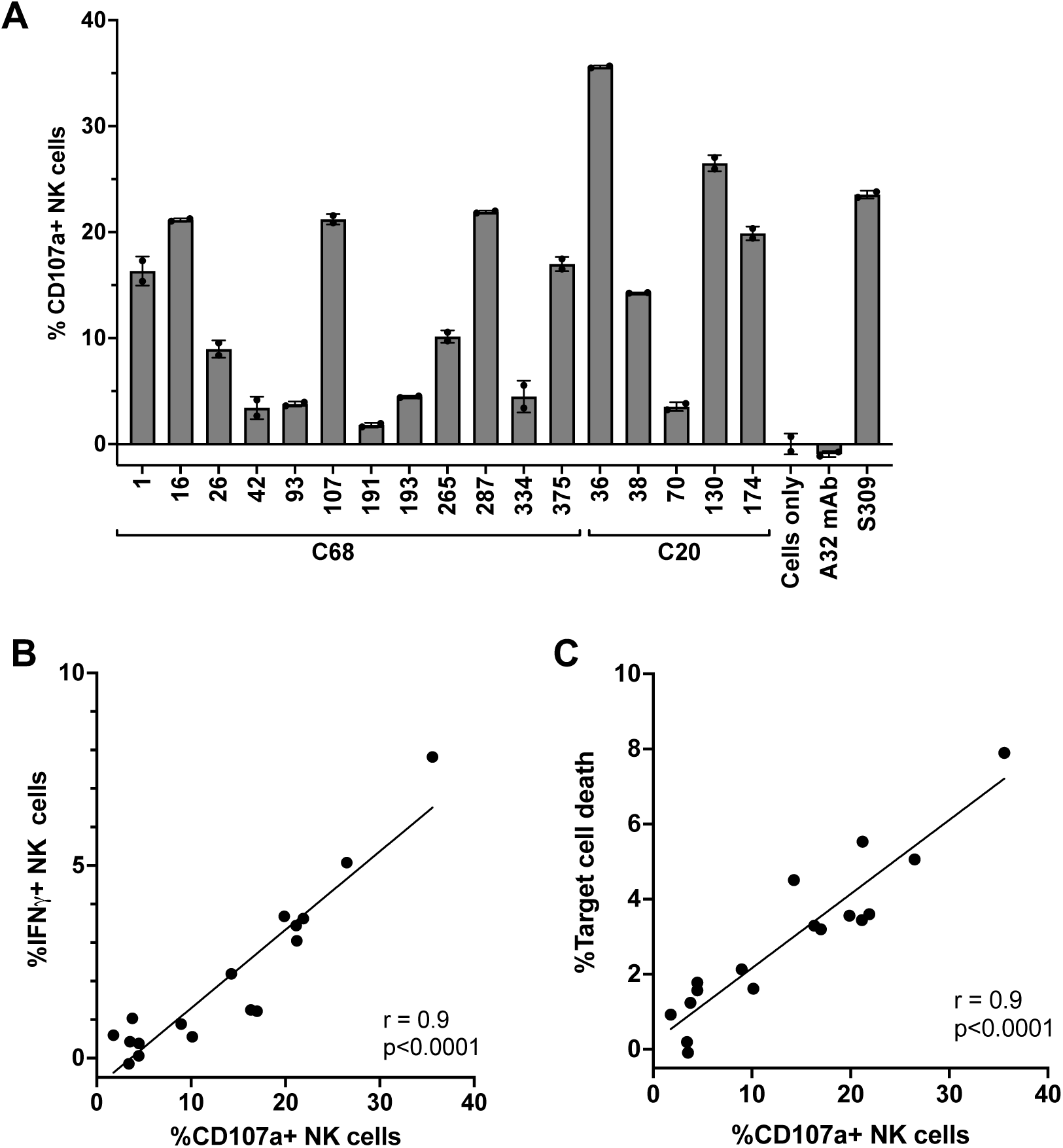
Antibody-dependent cellular cytotoxicity (ADCC) activity of S2 mAbs. S2 mAbs that were able to bind WH-1 spike trimer and exhibited breadth across SARS-CoV-2 VOCs (n=17) were evaluated for ADCC activity in a flow cytometry-based assay. mAbs (5 μg/mL) were incubated with target cells (CEM.NKr CCR5+ cells expressing GPF-tagged D614G spike) and mixed with effector cells (PBMCs) isolated from a healthy donor (at a 1:10 ratio) for four hours. Subsequently, target cell death and NK cell activation (CD107a+ and intracellular IFN-γ+) were measured. (A) Percentages of activated NK cells (CD107a+) are shown for C68 and C20 S2 mAbs, and controls HIV mAb A32, spike RBD mAb S309, and target cells and PBMCs without mAbs (“Cells only”). Values are the average of technical replicates from an experiment using PBMCs from a single donor (Donor 1) and the error bars indicate the standard deviation of the average. **(B, C)** Plots showing correlation between % IFN-γ+ NK cells **(C)** or % target cell death **(D)** with the % CD107a+ activated NK cells assessed by Pearson correlation. Data using PBMCs from the second donor are shown in **S6 Fig**.

### Binding competition groups reveal a diverse landscape of shared S2 epitope regions

Only a few epitopes of S2-specific antibodies have been characterized (66, 96), and the range of epitope targets within S2 is unclear. Therefore, we performed competition ELISAs to begin to define S2 antibody groups with similar epitopes using published S2 mAbs. These published mAbs with known epitopes were selected to represent the most commonly described S2 epitope regions: FP (COV2-2164 (79), 76E1 (28), C13B8 and VP12E7 (42)); HR1 (COV2-2002 and COV2-2333 (79)), and SH (B6 (70) and CV3-25 (54)), each mAb representing slightly different binding residues within the region or different angles of binding approach. All of the C68 and C20 mAbs that were able to bind WH-1 spike trimer at OD450nm ≥ 1.0 at 1 μg/mL (**Fig 1**) were included in the competitions, which represented 35 of the 40 S2 mAbs. Percent competition was determined for each mAb combination (**S7 Fig**). Using unsupervised clustering, we defined five major competition groups of S2 antibodies based on the clustering of the blocking mAbs (**Fig 5A**). Two groups, Group 1 and Group 4, included published S2 mAbs with known epitopes, whereas Groups 2 and 3 appeared to represent new S2 antibody groups as they did not compete with the published S2 mAbs. Group 5 was a collection of mAbs to diverse epitopes that largely did not compete with any other mAb tested.

**Fig 5.**
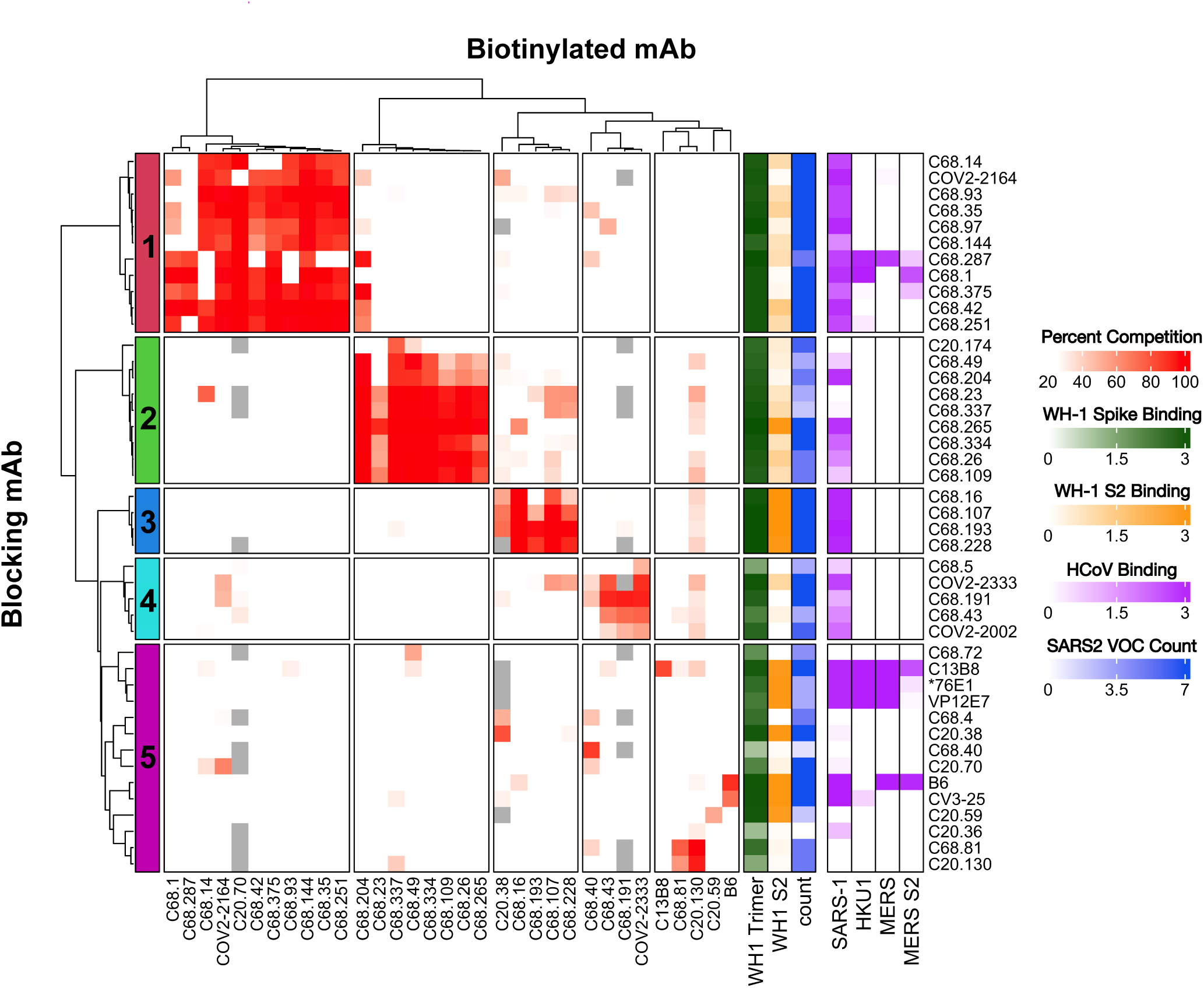
Antibody binding competition-defined sets of S2 mAbs with shared epitope regions and V(D)J gene usage. **(A)** Cross competition matrix of C20 and C68 S2 mAbs. Antibodies were selected based on binding to WH-1 prefusion-stabilized spike trimer (OD450nm > 1.0). Thirty-five of the forty mAbs were assessed. Several reference mAbs with known epitopes were also included in the competitions: FP (COV2-2164 (79), 76E1 (28), C13B8 and VP12E7 (42)); HR1 (COV2-2002 and COV2-2333 (79)), and SH (B6 (70) and CV3-25 (54)). Blocking mAbs (rows) were added at 2.5 μg/mL, followed by biotinylated mAbs (columns) at 100 ng/mL. All mAbs were tested in both directions as the blocking mAb and biotinylated mAbs. Any biotinylated mAb that did not reach at least 40% of the max OD450nm when competing with negative control mAb (HIV mAb VRC01) were removed from the analysis and are shown as grey boxes in the heat map. The heat map is colored based on the percent competition between any given pair of mAbs, which are averages of two technical replicates, with more red denoting higher competition. Any percent competition value <25% was set as white in the color scale in the heat map. Hierarchical clustering of the blocking mAbs identified five groups, marked on the left and separated into boxes. On the right of the matrix, gradients depict the binding features of each mAb to WH-1 spike trimer (green), WH-1 S2 protein (orange), spike trimers from SARS-CoV-1, HCoV-HKU1, MERS-CoV and MERS-CoV S2 protein (purple), and the count of SARS-CoV-2 VOC spike trimers bound by that mAb with at least half the maximal OD450nm to demonstrate SARS-CoV-2 breadth (blue).

We were able to predict the epitope region of two of the S2 epitope groups based on competition with published S2 mAbs. Group 1 was a large cluster consisting of ten C68 mAbs and one C20 mAb (**Fig 5**) that competed with the published antibody COV2-2164 (79), which was reported as an FP mAb due to binding to key residue 814 in the FP region. Group 1 mAbs and COV2-2164 bound prefusion-stabilized WH-1 spike trimer well in ELISAs at or near saturation (**Fig 1**). This group also had broad binding to SARS-CoV-2 VOCs; almost all of the Group 1 mAbs bound well to all seven of the SARS-CoV-2 spikes tested (OD450 > half maximal) and they bound to SARS-CoV-1 spike, as indicated by the colored annotations in the heatmap (**Fig 5**). The C68 mAbs that demonstrated cross-reactive binding to spike trimers from related HCoVs MERS-CoV and HCoV-HKU1, C68.1, C68.287, and C68.375, were also members of Group 1. Therefore, the Group 1 mAbs likely bind a highly conserved region that could include a portion of the FP that is accessible in the prefusion trimer. Based on competition with the published antibodies COV2-2002 and COV2-2333 (79), the four C68 mAbs in Group 4 likely bind the HR1 domain in S2 (912–984) (**Fig 5**). Similar to the binding profile of these published mAbs, the C68-derived Group 4 mAbs exhibited very weak binding to the S2 protein. These mAbs had reduced binding to the Omicron VOC spike trimers as well (**Fig 1**), which have key amino acid mutations at Q954 and N969 in HR1 compared to WH-1.

Groups 2 and 3 were two clusters of mAbs that did not compete with any of the published antibodies we tested (**Fig 5**). Group 2 was a large group of mAbs, eight from C68 and one from C20, with strong binding to the WH-1 spike trimer and weak binding to S2 protein. These mAbs tended to have broad but moderate binding to the SARS-CoV-2 VOCs and less binding to SARS-CoV-1 spike compared to Group 1 mAbs. C68.204 clustered with Group 2, but also competed strongly with several members of Group 1, specifically it was blocked by the Group 1 mAbs but it was not able to block those mAbs. Group 3 consisted of five mAbs, four from C68 and C20.38. All of these mAbs bound to saturation to WH-1 S2 peptide, all of the spike trimers from SARS-CoV-2 VOCs, and SARS-CoV1 indicating that they likely bind a conserved, highly accessibly epitope on the spike trimer. Finally, Group 5 was comprised of all the remaining mAbs that did not strongly compete with other mAbs and thus each likely represent unique epitope targets, with the exception of C20.130 and C68.81, which had strong cross competition between them (**Fig 5**). Notably, the published mAbs that bind to the SH region (B6 and CV3-25) competed with each other but did not cluster with any of the S2 mAbs we identified suggesting SH mAbs were not represented among the 40 mAbs isolated from C68 and C20. Further, the other published FP mAbs, C13B8, 76E1, VP12E7 did not compete with any of the C68/C20 mAbs, nor did they compete with each other suggesting there are other immunogenic regions in S2 that are commonly targeted by SARS-CoV-2 mAbs that have not been fully described.

We next examined whether mAbs in competition groups also had common V(D)J gene usage. As noted above, IGHV1-69, IGHV3-30, IGHV3-7 were commonly used genes in the mAbs we isolated and across reported SARS-CoV-2 mAbs (40, 50, 77–80). Here we observed enrichment of these genes in select competition groups: Group 1 mAbs, including the published mAb COV2-2164 (79), frequently used IGHV1-69 especially when combined with IGKV3-11, which has been observed as a public clonotype for S2 mAbs (82); Group 2 mAbs were enriched for the IGHV3-30 gene; and the majority of the mAbs in Group 3 used the IGHV3-7 gene (**S8 Fig**). The IGHV3-7 clonotype is seen in the HR1 published antibodies COV2-2002 and COV2-2333. COV2-2333 did compete moderately with Group 3 mAbs C68.107 and C68.228 when COV2-2333 was blocking (57% and 53% competition, respectively). This result indicates that there could be some shared or overlapping epitope between Group 3 and HR1 antibodies. Collectively, this group also tended to have longer CDR3 segments in comparison to other C68 and C20 mAbs. Overall, we were able to cluster the S2 mAbs into several competition groups, indicating shared epitope regions, and two of the groups competed with published antibodies in the FP (Group 1) and HR1 regions (Group 4) and another (Group 3) showed evidence for some HR1 features. However, many of the mAbs did not compete with published antibodies suggesting a diverse landscape of S2 epitopes that go beyond several of the commonly described regions.

### Linear epitope mapping identified C20.119 as an FP-neutralizing antibody

To date, most S2-specific neutralizing mAbs that have been identified target the FP (816–833) or SH regions (1141–1160) (28, 42, 43, 45, 51, 97). However, these regions can be challenging to map, especially FP, because the epitopes are often masked in prefusion-stabilized spike, which precludes the use of many standard epitope mapping methods, such as the binding competition we performed here. C20.119, which was the only neutralizing mAb in this set of S2 antibodies, bound well to the non-stabilized S2 protein but did not bind prefusion-stabilized trimer (**Fig 1**) and thus was not tested in our competition assays. Therefore, to determine if any of the mAbs identified here bind to linear epitopes in S2, we comprehensively mapped binding sites across spike using a phage-display immunoprecipitation sequencing (PhIP-seq) approach (98, 99). Previous work by our group and others has revealed that the FP and SH/HR2 regions can present as linear epitopes in individuals post-SARS-CoV-2 infection (28, 38, 52, 63, 99, 100), and these regions were detected as dominant responses in plasma from C20 and C68 using PhIP-seq (**S9 Fig**). Therefore, we employed two previously described phage-display libraries, both containing peptides spanning the complete extracellular domain of the WH-1 Spike protein, to identify SARS-CoV-2 peptides bound by the S2 mAbs.

First, C20 mAbs were screened with a library consisting of peptides 31 amino acids (aa) in length with 30 aa of overlap (98). Further, this library was designed for deep mutational scanning (DMS) enabling us to interrogate each aa at every position within the enriched peptides for reduced binding (98). Therefore, this library was termed “Phage-DMS” library. In technical replicates, we observed strong enrichment of a series of sequential peptides for two for the 11 C20 mAbs, C20.59 and C20.119 (**S10A Fig)**, neither of which were mapped by competition suggesting they represent additional epitope groups. The peptides for C20.59 spanned residues 790 to 810 in the WH-1 spike, which is just upstream of the FP region (**Fig 6A**). The peptides that were enriched for binding by C20.119 covered the region from 810 to 830 (**Fig 6B**), which includes FP and the S2’ cleavage site. None of the nine other C20 mAbs bound to linear peptides in this library (**S10A Fig**). By DMS, we observed loss of binding of C20.59 with numerous amino acid mutations to sites 794-804 (**Fig 6A**). Based on the ELISA binding data, C20.59 was able to bind spike trimer from WH-1 and Delta but lost the ability to bind spike trimers from Omicron VOCs (**Fig 1**), suggesting that its epitope is not well conserved across SARS-CoV-2 VOCs. The region bound by C20.59 contains a mutation that first emerged in Omicron BA.1 and is present in all subsequent VOCs at D796Y (**Fig 6C**), which was observed as an escape mutation in the Phage-DMS library profile as well (**Fig 6A**). The escape profile for C20.119 also refined the key binding residues to 813, 815-6, 818-20, and 822-5 (**Fig 6B**). Based on a sequence alignment of these residues across major SARS-CoV-2 VOCs over time, the C20.119 epitope, which is centered around the S2’ cleavage site, is completely conserved (**Fig 6C**).

**Fig 6.**
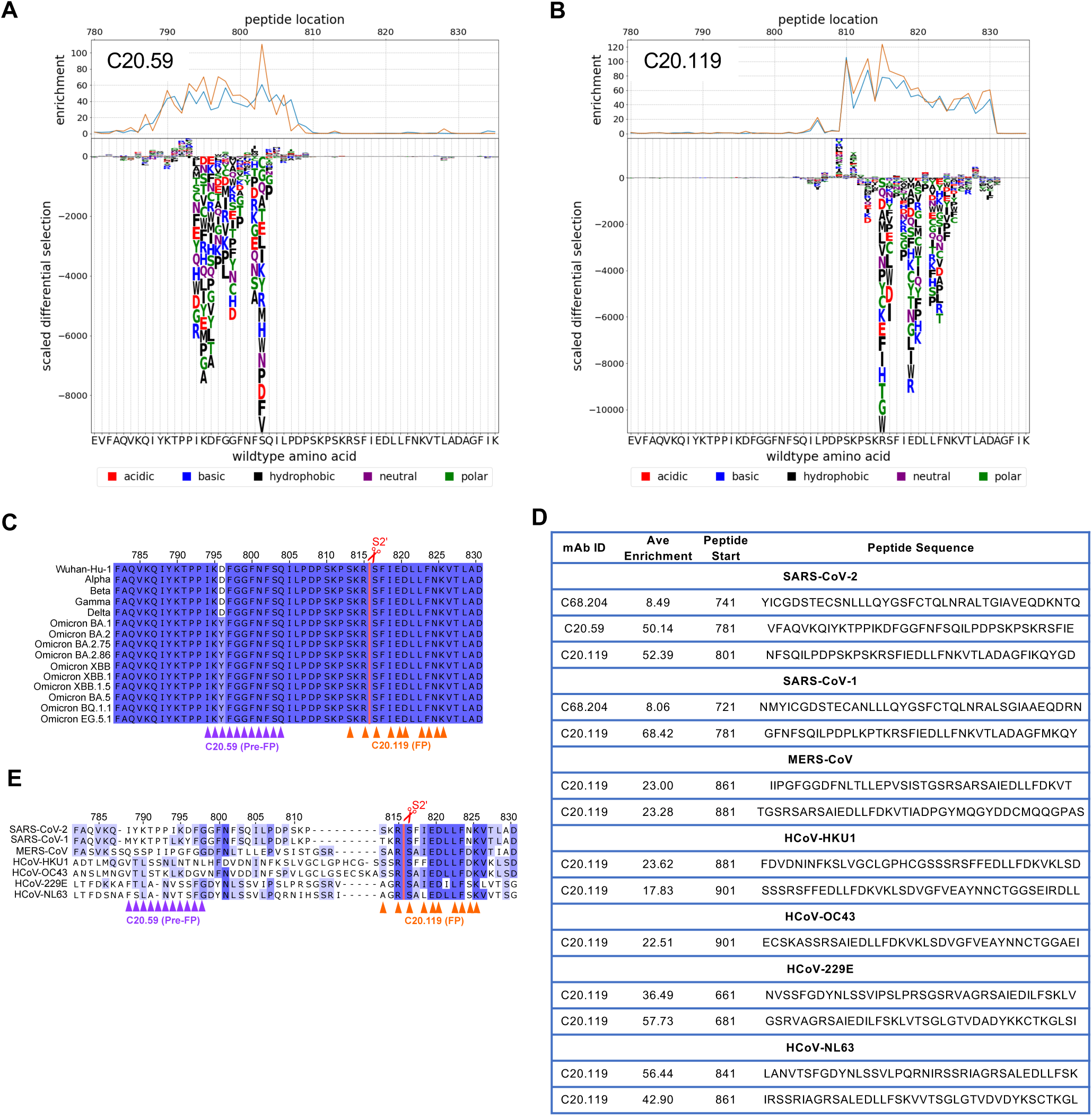
Epitope mapping by PhIP-seq identified linear epitopes in and around the FP region of S2 subdomain for mAbs C20.59, C20,119, and C68.204. **(A,B)** Each mAb was incubated with the phage-display spike peptide library, which consisted of 31 amino acid (aa) peptides with a 30 aa overlap (Phage-DMS library). Along with the Wuhan-Hu-1 wildtype sequences, each peptide was also represented with the central residue mutated to every possible amino acid to observe loss of mAb binding, or escape, with each mutation. The top graphs display the binding enrichment of peptides in that region in the WH-1 library for each mAb, with experimental replicates shown in the orange and blue lines for C20.59 **(A)** and C20.119 **(B)**. The bottom logo plots depict the effect of single mutations on binding of the mAb to the peptide. The height of the letter corresponds to the size of the effect that mutation had on the binding or the scaled differential enrichment with the mutation compared to the wildtype sequence. Mutations that resulted in a positive (>0) scaled differential enrichment are changes that increased mAb binding to the peptide, whereas a negative scaled enrichment is due to reduced binding with that mutation. The letter color signifies the chemical property of the amino acid. The WH-1 or wildtype sequence is shown on the x axis. **(C,D)** Sequence alignments across SARS-CoV-2 VOCs **(C)** or HCoVs **(E)** of the region upstream of and including FP (WH-1 aa 782-830). The key escape residues in the epitopes for C20.59 (purple) and C20.119 (orange) are noted under the alignment by the triangles. The color of the residues in the alignment denotes the percent identify at that site across the sequences shown. The S2’ cleavage is marked by the red line and scissors. **(D)** Peptides enriched in the pan-CoV library after binding with the three S2 mAbs C20.59, C20.119, and C68.204. The pan-CoV library included peptides covering the entire spike sequence from HCoVs SARS-CoV-2 WH-1, SARS-CoV-1, MERS-CoV, HCoV-HKU1, HCoV-OC43, HCoV-229E, and HCoV-NL63. Each mAb was run as technical replicates and the enrichment scores were averaged. Significantly enriched peptides (p<0.5) were identified and those peptides with an average enrichment greater than the off-target S1 regions are shown for C20.59, C20.119, and C68.204.

To map potential linear epitopes in the larger collection of C68 mAbs, we used a lower resolution library consisting of peptides 39 aa in length that overlapped by 19 aa and comprehensively covered the WH-1 spike sequence (99). This library also contained peptides covering spike protein sequences of closely related HCoVs, allowing us to examine cross reactivity of these mAbs. This library was called the “pan-CoV” library. C20.119 and C20.59 were also assessed with the pan-CoV library to compare results between the two libraries and assess HCoV breadth for these mAbs. As expected, we observed enrichment of peptides with C20.119 and C20.59 (**S10B Fig**), and the peptides bound in this lower resolution library covered the residues identified in the finer mapping performed with the Phage-DMS library (**S10A Fig & Fig 6D**). In technical replicates, most of the mAbs tested showed no peptide enrichment in S2 at p-value < 0.05 and for those peptides that were identified, many had relatively low enrichment scores, less than five as compared to >50 for C20.119 and C20.59 (**S10B Fig**). We also noted some mAbs had enriched peptides that mapped to the S1 subunit. Regions of S1 are proximal to S2 in the prefusion spike trimer (PDB 6VSB), so while it is possible a linear region in S1 could be contacted by an S2 mAb, the fact that none of the S2 mAbs described here bound S1 protein in ELISAs (**Fig 1**) suggested S1 regions are not a major part of the epitopes. Using the scores of the S1 peptides to set a threshold (score > S1 peptide enrichment average + 2SD), we identified one additional mAb, C68.204 that bound a peptide upstream of FP (741–779) (**Fig 6D & S10B Fig)**, which is consistent with the competition results where its binding to spike was blocked by Group 1 mAbs that we predict have an FP-involved epitope (**Fig 5**). None of the C20/C68 S2 mAbs bound to SH peptides in either library. We confirmed that published SH mAbs, CV3-25 (47) and 1A9 (101), could bind linear peptides in this region in our library (**S11 Fig**). Despite evidence SH is an apparent epitope in the antibody plasma response in these individuals based on our mapping (**S9 Fig**), these results suggest we did not isolate any mAbs that bound this region.

Given the sequence conservation of the S2 region across HCoVs, broad coronavirus reactivity of S2 mAbs has been described (39, 59). Therefore, we investigated if the three C20/C68 mAbs that bounds SARS-CoV-2 peptides exhibited binding to linear peptides in the pan-CoV library (99). C20.59 did not bind to any of the peptides in the other HCoV libraries at p-value < 0.05, which we predicted given this region is less well conserved between the HCoVs (**Fig 6E**). C68.204 bound to a peptide in the SARS-CoV-1 library (**Fig 6D**) that is in a region conserved between SARS-CoV-2 and SARS-CoV-1 and overlaps with the SARS-CoV-2 peptide identified (**S12 Fig**), which is consistent with the ability of this mAb to bind to the SARS-CoV-1 spike trimer (**Fig 1**). Finally, C20.119 bound peptides in all six of the HCoV libraries with high enrichment scores for all (**Fig 6D**). In the spike sequence alignment, we noted the peptides mapped to the same highly conserved sequence across all of the HCoVs at the S2’ cleavage site of SARS-CoV-2 (**Fig 6E**), which included the series of residues identified in the escape profiling with the higher resolution Phage-DMS library (**Fig 6B**). Overall, most of the S2 mAbs did not bind linear peptides; however, we were able to define the epitopes of three S2 mAbs, in or upstream of the FP region, one of which, C20.119, had broad binding across alpha and betacoronaviruses.

## DISCUSSION

In this study, we describe a collection of mAbs targeting diverse epitopes within the S2 subunit of the spike glycoprotein from two SARS-CoV-2 convalescent patients. To date, mAbs that target the more conserved S2 have been understudied as compared to those that bind S1 regions, such as RBD, which includes those antibodies that were previously authorized to treat or prevent infection with early strains of SARS-CoV-2 (11, 102–107). However, the RBD is one of the most variable regions across VOCs, which has led to immune escape, whereas the S2 region is highly conserved across SARS-CoV-2 VOCs. In the set of S2 antibodies presented here, we observed multiple shared epitope groups amongst the mAbs, many of which are likely to be conformational as they did not bind a comprehensive library of overlapping linear peptides that covered the S2 region. Functionally, we identified S2 mAbs with neutralizing or non-neutralizing (i.e., ADCC) activity, both of which have been suggested to be correlates of protection against SARS-CoV-2 (108) and demonstrate *in vivo* efficacy both as prophylactics and therapeutics in model systems (44, 51, 54, 56, 96, 109). Notably, many of the S2 mAbs displayed potent ADCC activity against SARS-CoV-2, with a subset having ADCC levels similar to S309, an RBD mAb that has dual neutralizing and Fc-effector functions, both contributing to its *in vivo* protective capabilities (49, 55).

The majority of previously described S2 antibodies target several common antigenic regions that are immunodominant and highly conserved in SARS-CoV-2: FP, HR1, HR2, and the SH region just upstream of HR2. Two of these regions, FP and SH, have been shown to be targets for neutralizing S2 mAbs (28, 42–48), but they can be challenging to map using methods that rely on binding to prefusion-stabilized spike both because the prefusion state can mask these regions and some of the introduced mutations to force stabilization are in these key epitopes (110). These regions can be detected as linear epitopes in serum after SARS-CoV-2 infection or vaccination (38, 99, 111, 112), and we noted significant enrichment of binding to linear peptides covering these regions in the serum of C20 and C68 as well. Some monoclonal S2 antibodies targeting these regions also bind linear epitopes (45, 51, 96). Therefore, to define epitope classes for the large collection of S2 mAbs described here, we utilized two complementary methods, phage display with deep sequencing and binding competition mapping with known S2 mAbs, which allowed us to either identify linear epitopes or cluster mAbs that appeared to share epitopes. With phage display and deep sequencing, we were able to resolve discrete epitopes for three of the mAbs all to the Pre-FP/FP region, and one mAb, C20.119, targeted a region that contains the S2’ cleavage site. However, we did not identify any S2 mAbs that mapped to SH regions, nor did we observe mAbs that could compete with known SH mAbs, suggesting that the S2 epitopes represented in this collection of mAbs might span beyond those commonly described.

In addition to the FP mAbs identified with phage display sequencing, by binding competition mapping we defined four groups that cross compete within each group indicative of a shared epitope region. A fifth group clustered together not because of cross-competition, but rather the lack of competition with other larger groups of mAbs, suggesting these mAbs bind unique or rare epitopes, or that the angle of binding does not lead to competition. For the defined classes with cross-competition, two groups competed with a published S2 mAb with a known epitope. Three C68 mAbs competed with the HR1 antibodies, COV2-2333 and COV2-2002 (79). A large group of mAbs (Group 1) that included those with binding breadth to other HCoVs competed with published antibody COV2-2164 (79), which binds to residues in the highly conserved FP region. However, we saw no competition between the antibodies in this group and the other published FP antibodies we tested (76E1 (28), C13B8 and VP12E7 (42)). We did observe competition between Group 1 mAbs and the Group 2 mAb C68.204, which we determined binds a peptide upstream of FP, further supporting that these two large groups of S2 mAbs bind epitopes in Pre-FP/FP. None of the other mAbs in competition Groups 1 or 2 bound a linear peptide, which suggests the epitopes in this region are largely conformational and accessible in prefusion spike. Two groups of mAbs did not compete with any of the published antibodies and currently have unknown epitopes, a set of four in Group 3 that cross-competed and another eight in Group 5 that did not compete with any other mAb. These mAbs did not bind linear peptides in the phage display, again indicating that these mAbs are binding to conformational epitopes. Together, based on four epitope groups identified by competition binding, two distinct epitopes for 59 and 119 and lack of SH mAbs, our results suggest at minimum eight epitopes in S2, but certainly more based on eight with no cross competition. Thus, altogether the mapping data indicate broad responses to diverse targets across S2 including a range of epitopes not presently described.

We have identified a mAb, C20.119, that targets a linear epitope in the FP region and has broad but only moderately potent neutralizing activity against SARS-CoV-2 VOCs including the more recent Omicron VOCs XBB.1.5 and BQ.1.1. Compared to other reported FP mAbs, C20.119 has similar potency to 76E1 from Sun, et al. (28), and is roughly 10 to 20-fold more potent than several mAbs from other studies (42, 43, 113) against WH-1. Notably, several of these mAbs demonstrated *in vivo* protection, as prophylactics and treatments, in animal models challenged with SARS-CoV-2 virus. All of the FP mAbs described appear to inhibit membrane fusion by binding this region (42, 43, 50), so we predict C20.119 would work in a similar fashion. While there are no mutations in the epitope of C20.119 in the Omicron VOCs, we did observe a reduction in potency against the Omicron pseudoviruses as compared to early strains WH-1 and Delta. A possible explanation for this finding is that Omicron variants preferentially utilize endocytosis over membrane fusion for initial target cell entry, the latter of which is preferred by WH-1 and Delta (114), thereby attenuating the potential neutralizing effect of an FP-targeting mAb. This could also reflect differences in escape based on the viral background used for phage display DMS as we have seen with other mAbs (68). Consistent with this, mutations throughout other regions of spike in Omicron VOCs have been shown to alter the structure of the trimer and potentially the accessibility of the FP (115, 116).

Collectively, many of the S2 mAbs described here with functional activity and breadth against SARS-CoV-2 VOCs also exhibited cross reactivity to other related viruses including HCoVs and sarbecoviruses. The majority of the mAbs from C68 and three of the C20 mAbs, were able to bind SARS-CoV-1 spike trimer by ELISA. Cross-reactive SARS-CoV-1/SARS-CoV-2 antibodies are well described, as expected given that the S2 amino acid sequence identity between the two viruses is ∼90% (101, 117). Further, three of the C68 mAbs, were able to bind recombinant spike proteins from other betacoronaviruses MERS-CoV and/or HCoV-HKU1. These three mAbs also exhibited robust NK cell activation through their Fc-receptor functions, making them attractive candidates to further evaluate their efficacy against HCoVs. While the percent spike sequence similarity drops considerably between WH-1 and these HCoVs (97% with SARS-CoV-1 spike to 30% with HCoV-HKU1 (118)); several regions of S2, notably FP and the S2’ cleave site are highly conserved suggesting they are subject to functional constraint (118, 119). Indeed, our comprehensive linear epitope mapping revealed that C20.119, the neutralizing FP mAb, was able to bind linear peptides across all seven HCoVs. Further, C20.119 was able to neutralize pseudoviruses representing closely related, zoonotic sarbecoviruses that can utilize ACE2 for cell entry and have highly conserved S2 regions when compared to WH-1 (>90-100% in different regions of S2) (17). As is evident from SARS-CoV-1 and SARS-CoV-2, zoonotic spillover of these sarbecoviruses is a concern, so mAbs, such as C20.119, that can maintain function across these viruses are critical for preparing for these potential future spillover events.

The combined breadth and antiviral functions of S2 antibodies suggest they are an important component of an immune response most poised to protect against SARS-CoV-2 VOCS and HCoVs. As we have described here, the diversity of epitopes in the S2 response spans beyond what has been described and understanding the array of S2 epitopes can provide targets for vaccine design. Several groups have designed S2-focused vaccines in mice and have demonstrated that this strategy can elicit broad, cross-reactive antibodies that are protective against both SARS-CoV-2 and other HCoVs (36, 120, 121). Importantly, in mice that have a history of vaccination with the WH-1 spike, which similar to the human population that has seen several rounds of SARS-CoV-2 vaccines containing WH-1 spike, additional vaccination with an S2 only vaccine was able to induce a broader response as compared to additional WH-1 spike boosters (36). These studies support the incorporation of an S2-focused immunization strategy that could broaden the immune response priming it for future, diverse exposures. The antibodies we describe here provide additional information about the S2-targeted antibody response to SARS-CoV-2 and can contribute to the efforts to develop such novel vaccines.

## MATERIALS AND METHODS

### Study Participant and Specimens

Peripheral blood mononuclear cells (PBMCs) were collected 30-days post symptom onset (pso) from two individuals following SARS-CoV-2 infections who were enrolled in the Hospitalized or Ambulatory Adults with Respiratory Viral Infections (HAARVI) research study (122). Written, informed consent was obtained from individuals C20 and C68 at the time of enrollment in March 2020 and July 2021, respectively. The study was approved by Institutional Review Boards at University of Washington and Fred Hutchinson Cancer Center.

### Single-cell sorting for spike-specific memory B cells

Memory B cells expressing receptors encoding SARS-CoV-2 spike-specific antibodies were sorted from 30-day pso PBMCs using standard methods to identify memory B cells that bind the spike antigen (68, 123, 124). PBMCs were stained using a mouse anti-human antibody cocktail to cell surface markers: CD3-BV711 (BD Biosciences, clone UCHT1), CD14-BV711 (BD Biosciences, clone MφP9), CD16-BV711 (BD Biosciences, clone 3G8), CD19-BV510 (BD Biosciences, clone SJ25C1), IgM-FITC (BD Biosciences, clone G20-127), IgD-FITC (BD Biosciences, clone IA6-2). A baiting approach was used to isolate the spike-specific B cells: C20 PBMCs were baited with APC/PE-labeled WH-1 spike S2 protein (Acro Biosystems, cat. S2N-C52E8) and C68 PBMCs with a pool of APC/PE-labeled Delta HexaPro spike trimer protein (obtained from David Veesler) and WH-1 spike S2 protein (Acro Biosystems, cat. S2N-C52E8). The gating strategy used for the C20 B-cell sort is shown in the **S13 Fig** and the C68 B-cell sort is shown in (68).

### Reconstruction of antibodies

Antibody gene sequences were recovered from the individually sorted B cells with gene-specific primers and sequenced using a previously optimized pipeline (68, 123–128). The variable region sequences were assessed with the IMGT database (129) to determine if they are productive, in-frame variable regions. Those sequences that did not have an intact open reading frame (ORF) were excluded. Productive heavy and light variable regions paired in the same well (i.e., same B cell), were cloned into the appropriate IgG1 gamma, kappa, or lambda constructs and transfected into FreeStyle™ 293-F Cells (Invitrogen) to produce full-length mAbs for functional characterization as previously described (68, 123–128). For C20, 251 B cells were isolated into 96-well plates, and paired and productive (i.e., intact ORF) heavy and light chain sequences were isolated from 31 B cell wells. Twenty-six of the 31 heavy and light chain pairs were able to product sufficient antibody for characterization. Eleven of the antibodies bound to SARS-CoV-2 WH-1 spike trimer or S2 protein in direct ELISAs above background (determined by absorbance at OD450nm greater than the HIV mAb VRC01 OD450nm of + 3 standard deviations). For C68, 384 B cells were isolated into four 96-well plates, and 118 B cell wells had paired heavy and light chain sequences that produced sufficient antibody after transfection. Of those 118 mAbs, 29 were determined to be S2 specific based on binding to WH-1 spike trimer or S2 protein above background as described above. The remaining mAbs bound to other spike subunits (45 RBD, 17 NTD, 3 CTD) or were not spike specific (n=24).

### Inference of germline genes and sequence annotation with partis

Sequences were annotated with V, D, J genes, deletion lengths, and non-templated insertions and grouped into clonal families using partis (75). This includes automatic germline inference (74) for detecting previously unknown alleles. Partis groups together sequences stemming from a single rearrangement event with a multi-step process: first grouping together observed sequences with similar inferred naïve sequences, then applying a more accurate (but slower) likelihood-based method for more difficult cases (76). Since each well contained a single cell, reads from each well were treated as if from the same 10X-style droplet for input to partis’s paired clustering, which leverages simultaneous heavy and light chain clonal information to improve the clustering accuracy of both chains (75). Data analysis and visualization of mAb gene features was performed plotted using GraphPad Prism (v10). Gene information for the mAbs including variable region aa sequences, inferred germline, percent somatic hypermutation, and CDR3 length for the 40 S2 mAbs in this study are shown in **S1 Table**.

### Binding determined by direct enzyme-linked immunosorbent assays (ELISAs)

Binding of the mAbs to recombinant spike trimers and spike subunit proteins was assessed by direct ELISAs as previously described (68). The recombinant spike trimers used in these assays were as follows: Wuhan-Hu-1 (Sino Biological, cat. 40589-V08H4), Delta (Sino Biological, cat. 40589-V08H10), Omicron BA.1 (Sino Biological, cat. 40589-V08H26), Omicron BA.2 (Sino Biological, cat. 40589-V08H28), Omicron BA.4/BA.5 (Sino Biological, cat. 40589-V08H32), Omicron XBB (Sino Biological, cat. 40589-V08H40), Omicron BQ.1.1 (Sino Biological, cat. 40589-V08H41), SARS-CoV-1 (Acro Biosystems, cat. SPN-S52Ht), MERS-CoV (Acro Biosystems, cat. SPN-M52H4), HCoV-HKU1 (Acro Biosystems, cat.SPN-H52H5). The spike subdomain proteins were SARS-CoV-2 S1 (Sino Biological, cat. 40591-V08H), SARS-CoV-2 S2 (insect cell derived, Sino Biologics, cat. 40590-V08B), SARS-CoV-2 S2 (mammalian cell derived, Sino Biologics, cat. 40590-V08H1), MERS-CoV S2 (Sino Biologics, cat. Cat: 40070-V08B). Data were analyzed and plotted using GraphPad Prism (v10).

### Epitope groups defined by competition ELISAs

All mAbs that bound to SARS-CoV-2 WH-1 spike trimer in direct ELISAs at OD450nm > 1.0 (35 of the 40 mAbs) were assessed in the competition ELISAs. Published mAbs with known epitopes were included to represent the most commonly described S2 epitope regions: FP (COV2-2164 (79), 76E1 (28), C13B8 and VP12E7 (42)); HR1 (COV2-2002 and COV2-2333 (79)), and SH (B6 (70) and CV3-25 (54)). The competition ELISAs were performed using a similar protocol to the direct ELISAs and described previously (68), with some alterations. The WH-1 spike trimer (Sino Biological, cat. 40589-V08H4) was used as the antigen in all ELISAs at a concentration of 1μg/mL overnight at 4°C and blocked with 3% BSA in wash buffer for one hour at 37°C. After washes, the non-biotinylated (“blocking”) mAb was applied at 2.5 μg/mL and incubated at 37°C for 15 minutes. Then, the biotinylated mAb was added at 100 ng/mL and incubated at 37°C for 45 minutes. The remaining washes, incubation with secondary antibody (anti-streptavidin IgG-HRP), and reaction with the TMB chromogen substrate to allow quantification of biotinylated mAb binding at absorbance OD450 was completed as previously described (68). Each of the mAbs was tested as the blocking mAb and biotinylated mAb in every combination. As a control, each biotinylated mAb was also incubated after blocking with the non-spike, HIV mAb VRC01, which should not compete with the spike-specific mAbs. Any biotinylated mAb that had an OD450nm <40% of the maximum OD450nm when VRC01 was blocking was excluded from the analysis. However, these mAbs were kept as blocking antibodies in the analysis. The OD450nm values were background-subtracted using the buffer-only wells, and the percent competition for each blocking/biotinylated mAb combination was calculated [(1-OD450_well_/average OD450_ave_ of VRC01 wells)*100]. Percent competition values were averaged across two technical replicates and any negative values were set to zero. The matrix of percent competition values for all blocking/biotinylated mAb combinations is shown in **S6 Fig**. Competition groups were determined using a hierarchical clustering algorithm utilizing a Pearson distance matrix and Ward’s Method, implemented as ward.D2 in the R environment (130). Group number was set as five for both rows and columns. Heatmaps and determination of competition groups were made using the ComplexHeatmap Package in the R environment (https://jokergoo.github.io/ComplexHeatmap-reference/book/index.html). A percent competition visual cutoff was also implemented in the heatmap where values <25% were set to the same color as 0%.

### Spike-pseudotyped lentivirus neutralization

Spike-pseudotyped lentiviruses were produced and titered as previously described (52, 68, 83). Codon-optimized plasmids with spike genes from SARS-CoV-2 strains (listed in **Fig 3** and generously provided by Amit Sharma, Jesse Bloom, and Marceline Côté), SARS-CoV-1 (Urbani strain, GenBank AY278741, from Marceline Côté), and zoonotic sarbecoviruses RsSHC014 and WIV1 (86) (a kind gift from David Veesler) and Pangolin-GD (131) (gifted from Pamela Bjorkman), were individually transfected with lentiviral helper plasmids into HEK293T cells for virus production. Neutralization of the pseudoviruses by the mAbs was performed as previously described (68). HEK293T-ACE2 cells expressing high levels of ACE2 (83, 132) were used as the target cells to measure infection and neutralization. A non-neutralizing antibody (e.g., HIV mAb VRC01) was run in parallel as a control. For each run, the average raw relative light units (RLUs) of the virus plus cells alone wells was compared to the average RLUs of the negative control mAb wells. The average of the virus plus cells wells was used to determine the maximum infectivity value as is standard practice; however, in runs where the virus plus cells RLU average was >1.3 fold higher than the negative control mAb average, the negative control average was used. Background signal from negative control wells (no mAb, no virus) was subtracted, replicate experiments were averaged, and the fraction of infectivity was calculated. For screening the pseudovirus neutralization potency of the S2 mAbs, percent neutralization at 10 μg/mL was calculated and for C20.119, two-fold mAb dilution curves were assessed, and the half maximal inhibitory concentrations (IC50s) were calculated with a non-linear regression fit for inhibition versus response constraining the bottom to 0, the top to 1, and HillSlope < 0. Data were analyzed and plotted using GraphPad Prism (v10).

### Antibody binding to cell-surface expressed SARS-CoV-2 spike protein

Binding of S2 mAbs to natively expressed SARS-CoV-2 spike protein on cells (kindly provided by Andrés Finzi) was assessed by flow cytometry. Briefly, 10^5^ CEM.NKr CCR5+ cells stably expressing GFP-tagged D614G spike protein (92) were incubated with mAbs at 5 μg/mL for 30 minutes at 37°C (consistent with the ADCC assay conditions, see below). Cells were washed twice, and stained with anti-human IgG, Fcγ fragment-specific PE (Jackson Immuno, cat. 109-115-098) for 30 min. Cells were then washed and fixed before acquisition on a Fortessa LSRII instrument (BD Biosciences) with BD FACSDiva software. Data was analyzed using FlowJo v10.7.1 (TreeStar). Positive (CV3-25 GASDALIE) and negative (HIV-specific A32) control antibodies were included in each experiment. An example gating strategy is shown in **S3 Fig**.

### SARS-CoV-2 Spike ADCC assay to detect activated NK cells

Cryopreserved PBMCs from healthy adult donors, used as effector cells in the assay, were thawed and allowed to rest overnight in RPMI supplemented with 10% FBS. Effector cells were mixed with target cells (CEM.NKr CCR5+ cells stably expressing GFP-tagged D614G Spike protein (92); provided by Andrés Finzi) at a 10:1 ratio, in the presence of S2 mAbs (5 μg/mL), protein transport inhibitor cocktail (1:500, eBioscience) and anti-CD107a APC (clone H4A3, BioLegend). Antibody-dependent NK cell activation was allowed to occur for 4 hours at 37°C and 5% CO_2_. The cells were then washed twice and stained for viability (Fixable Viability Dye eFluor 780, eBioscience) and with anti-CD56 BV605 (clone 5.1H11, Biolegend) and anti-CD3-BV711 (clone UCHT1, BD Bioscience) for 30 minutes at 4°C. The cells were then washed twice, fixed/permeabilized using IC Fix/Perm (Invitrogen) and stained with anti-IFNγ–PE (clone 4 S.B3, BioLegend) for 30 minutes at room temperature. Data was acquired on a Fortessa LSRII instrument (BD Biosciences) with BD FACSDiva software and analyzed using FlowJo v10.7.1 (TreeStar). Target cells were identified according to cell morphology by light-scatter parameters and expression of GFP and were assessed for viability. NK cells were identified by gating on lymphocytes by light-scatter parameters, exclusion of doublets and dead cells, and were defined as CD56+ CD3-. Gating strategies are shown in **S4 Fig**. Each assay was performed in technical duplicates and repeated with two independent effector cell donors. Positive (S309) and negative (HIV-specific A32) control antibodies were included in each experiment.

### Phage-display immunoprecipitation and deep sequencing

Linear epitope mapping of S2 mAbs and human plasma by phage-display immunoprecipitation sequencing (PhIP-seq) was performed as previously described (38, 52, 98–100). The first, higher resolution peptide library (termed “Phage-DMS” library) was designed from sequence from the Wuhan Hu-1 strain (GenBank:MN908947) covering the ectodomain of the spike protein (aa 1-1211), excluding the transmembrane and cytoplasmic domains. The library included peptides 31 aa long tiled every 30 aa for single residue resolution, and details on the design have been previously described (52). In addition, this library included variations of every wildtype peptide where each mutant peptide contained a single variable residue at the central position of the peptide; thereby resulting in 20 peptide sequences containing all possible mutations at each position along the protein. The second, lower resolution library (termed “pan-CoV” library) included peptides from 17 spike coding sequences: OC43-SC0776 (MN310478), OC43-12689/2012 (KF923902), OC43-98204/1998 (KF530069), 229E-SC0865 (MN306046), 229E-0349 (JX503060), 229E-932-72/1993 (KF514432), NL63-ChinaGD01 (MK334046), NL63-Kilifi_HH-5709_19-May-2010 (MG428699), NL63-012-31/2001 (KF530105), NL63-911-56/1991 (KF530107), HKU1-SI17244 (MH90245), HKU1-N13 genotype A (DQ415909), HKU1-Caen1 (HM034837), MERS-KFMC-4 (KT121575), SARS-Urbani (AY278741), SARS-CoV-2-Wuhan-Hu-1 (MN908947), bat-SL-CoVZC45 (MG772933). A single HIV-1 envelope sequence was also included for controls (BG505.W6.C2,DQ208458). Each peptide in this library was 39 aa in length and tiled with 19 aa overlaps, design previously described (99). Immunoprecipitation of S2 mAbs or plasma with both phage display libraries was performed as previously described (52, 99). In brief, deep 96-well plates were blocked with 3% BSA in Tris-buffered saline with 0.01% Tween (TBST) overnight at 4°C. The phage library was diluted (200,000 pfu/mL per unique peptide) and 1 mL of the diluted phage was added to each well. mAbs (10 ng per well) or plasma (assuming plasma IgG concentrations to be 10 μg/μL (133) added at 10 μg per well). Each mAb was run in technical replicates. After allowing the mAbs to bind overnight at 4°C rocking, the phage:antibody complexes were immunoprecipitated using 40 μL of a 1:1 mixture of protein A and protein G magnetic Dynabeads (Invitrogen) (4°C for 4 hours, rocking). The magnetic beads were separated and washed three times and resuspended in 40 μL of water. The bound phage were lysed and stored at −20°C until Illumina library preparation.

Illumina library preparation was performed as previously described (52, 98, 99). Phage DNA underwent two rounds of PCR to produce Illumina libraries containing adaptor sequences and barcodes for multiplexing. PCR products were cleaned with AMPure XP beads (Beckman Coulter) and quantified by Quant-iT PicoGreen according to manufacturer instructions (Thermo Fisher). Samples were pooled and deep sequenced on an Illumina MiSeq with 1×125 bp single end reads using a custom sequencing primer.

#### Phage sequence alignment and data analysis

Processing of sequence reads from the phage display protocols and downstream analyses of peptide enrichments are performed with the *phippery* software suite (134). In brief, *phippery* aligned raw reads to the corresponding Phage-DMS or pan-CoV peptide library reference via *Bowtie2* (135) and *Samtools* (136). Peptide enrichment was calculated as the observed relative abundance in a sample over that observed in the input peptide library. For peptides from the Phage-DMS library, differential selection was calculated as the log-fold-change of the mutant peptide enrichment versus the wildtype peptide enrichment. Scaled differential selection was defined as differential selection multiplied by the average wildtype enrichment at the locus of interest and its two adjacent loci; the use of averaging reduces artificially enhanced values due to noise. For peptides from the pan-CoV library, *p*-values of the enrichments were computed from the method described in (137), and accounted for multiple hypothesis testing; a peptide enrichment was considered significant if p < 0.05 in both sample replicates. Pan-CoV peptides that were enriched with a p < 0.05 in both replicates were considered potential binding peptides for that mAb. The cutoff of p < 0.05 was well calibrated by checking for the enrichment of HIV-1 envelope peptides; 4.8% of the HIV-1 peptides had p < 0.05.

## ACKNOWLEDGEMENTS

We thank Marceline Côté (University of Ottawa), Amit Sharma (Ohio State University), Jesse Bloom (Fred Hutchinson Cancer Center), David Veesler (University of Washington) and Pamela Bjorkman (Caltech) for generously providing spike plasmids for production of spike-pseudotyped lentiviruses. We also thank David Veesler (University of Washington) for providing Delta spike trimer to use as bait to capture Spike-specific B-cells and Andrés Finzi (Université de Montréal) for the CEM.NKr CCR5+ cells. We also would like to thank the participants and the study staff of the Hospitalized or Ambulatory Adults with Respiratory Viral Infections (HAARVI) study.

## Notes

### Competing Interest Statement

J.O. is a consultant for Aerium Therapeutics, Inc. H.Y.C reported consulting with Ellume, Merck, Abbvie, Pfizer, Medscape, Vindico, and the Bill and Melinda Gates Foundation. She has received research funding from Gates Ventures, and support and reagents from Ellume and Cepheid outside of the submitted work. J.O. and J.G are listed on a patent application (22-173-US-PSP2) and license agreement with Aerium Therapeutics, Inc for SARS-CoV-2 mAbs not described in this manuscript.

