## Supplemental Information for "The S2 subunit of spike encodes diverse targets for functional antibody responses to SARS-CoV-2"

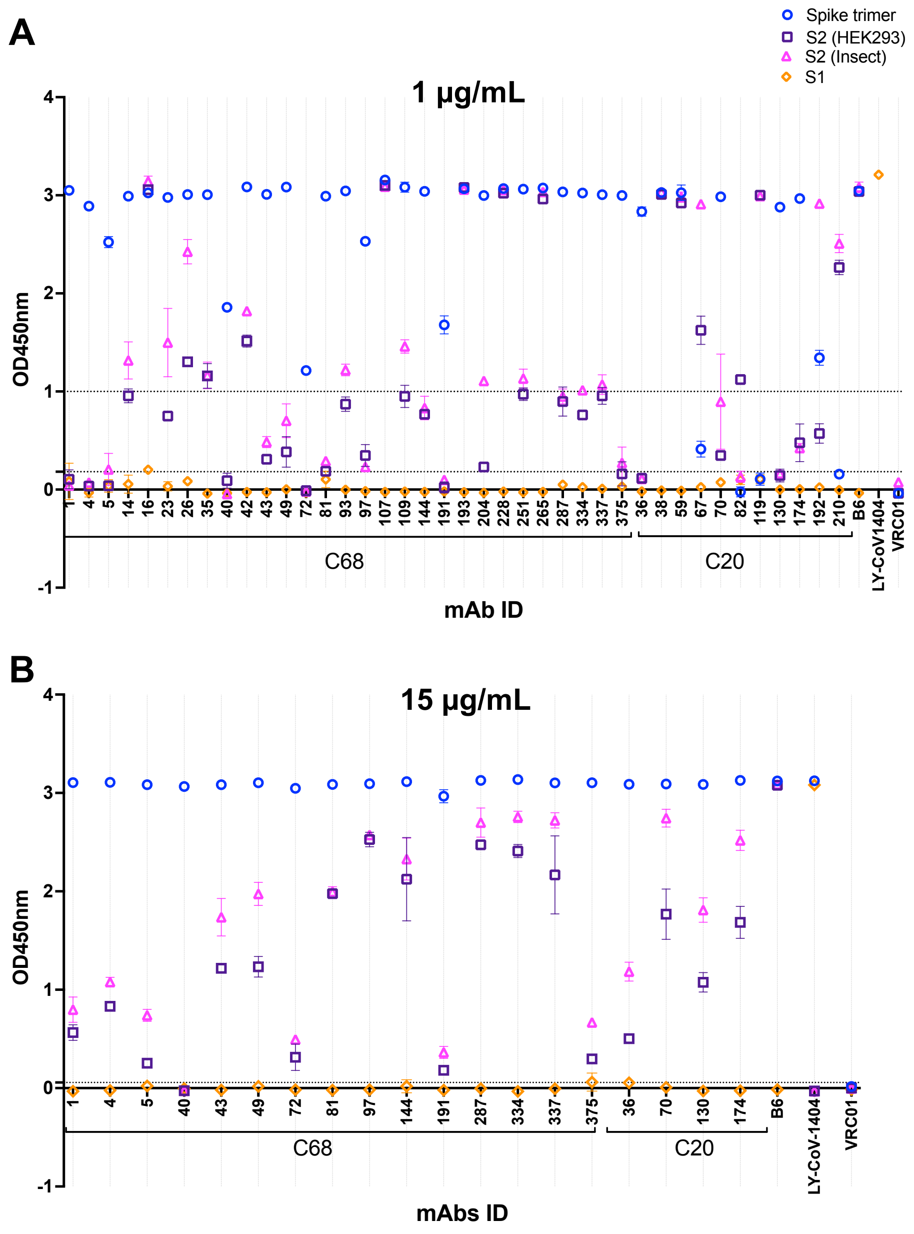
A
Spike trimer
S2 (HEK293)
S2 (Insect)
S1
1 μg/mL
OD450nm
mAb ID
C68
C20
1 4 5 14 16 23 26 35 40 42 43 49 72 81 93 97 107 109 144 191 193 204 228 251 265 287 334 337 375 36 38 59 67 70 82 119 130 174 192 210 B6 LY-CoV1404 VRC01
B
15 μg/mL
OD450nm
mAbs ID
C68
C20
1 4 5 40 43 49 72 81 97 144 191 287 334 337 375 36 70 130 174 B6 LY-CoV-1404 VRC01

S1 Fig.

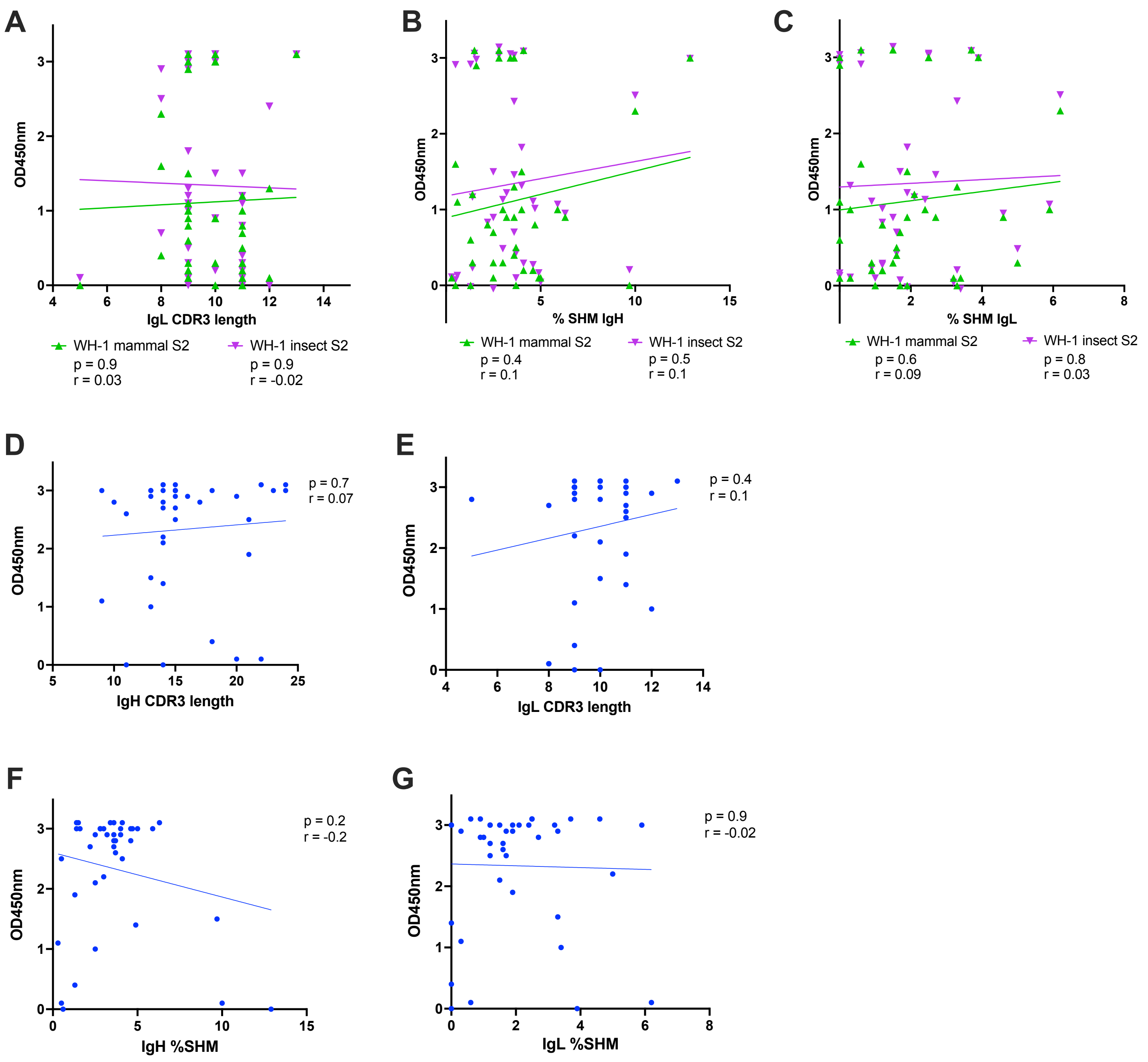
A
OD450nm
IgL CDR3 length
WH-1 mammal S2
p = 0.9
r = 0.03
WH-1 insect S2
p = 0.9
r = -0.02
B
OD450nm
% SHM IgH
WH-1 mammal S2
p = 0.4
r = 0.1
WH-1 insect S2
p = 0.5
r = 0.1
C
OD450nm
% SHM IgL
WH-1 mammal S2
p = 0.6
r = 0.09
WH-1 insect S2
p = 0.8
r = 0.03
D
OD450nm
IgH CDR3 length
p = 0.7
r = 0.07
E
OD450nm
IgL CDR3 length
p = 0.4
r = 0.1
F
OD450nm
IgH %SHM
p = 0.2
r = -0.2
G
OD450nm
IgL %SHM
p = 0.9
r = -0.02

**S2 Fig.**

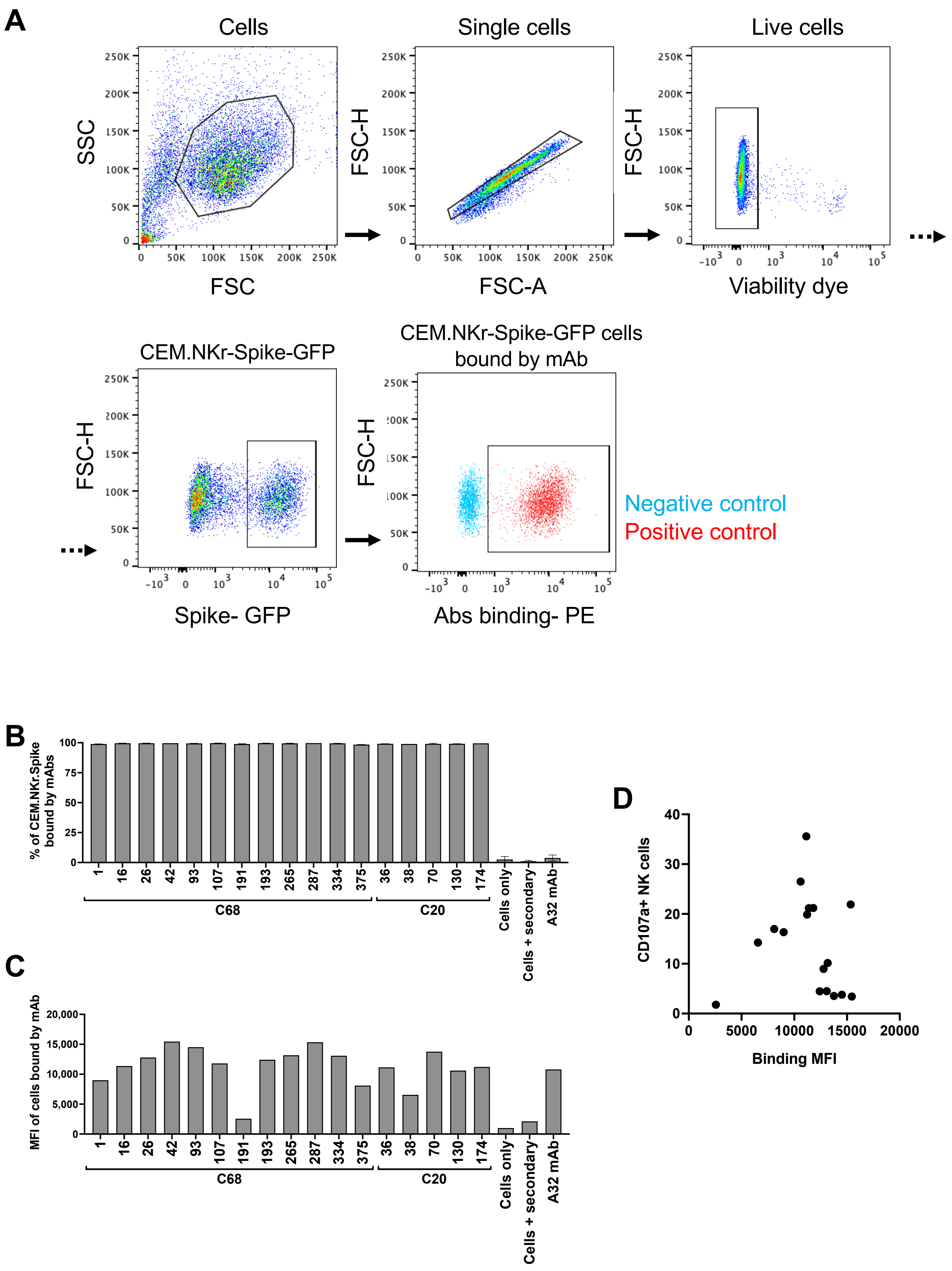
A
Cells
Single cells
Live cells
SSC
FSC
FSC-H
FSC-A
Viability dye
CEM.NKr-Spike-GFP
CEM.NKr-Spike-GFP cells bound by mAb
Spike- GFP
Abs binding- PE
Negative control
Positive control
B
% of CEM.NKr.Spike bound by mAbs
C68
C20
Cells only
Cells + secondary
A32 mAb
C
MFI of cells bound by mAb
D
CD107a+ NK cells
Binding MFI

S3 Fig.

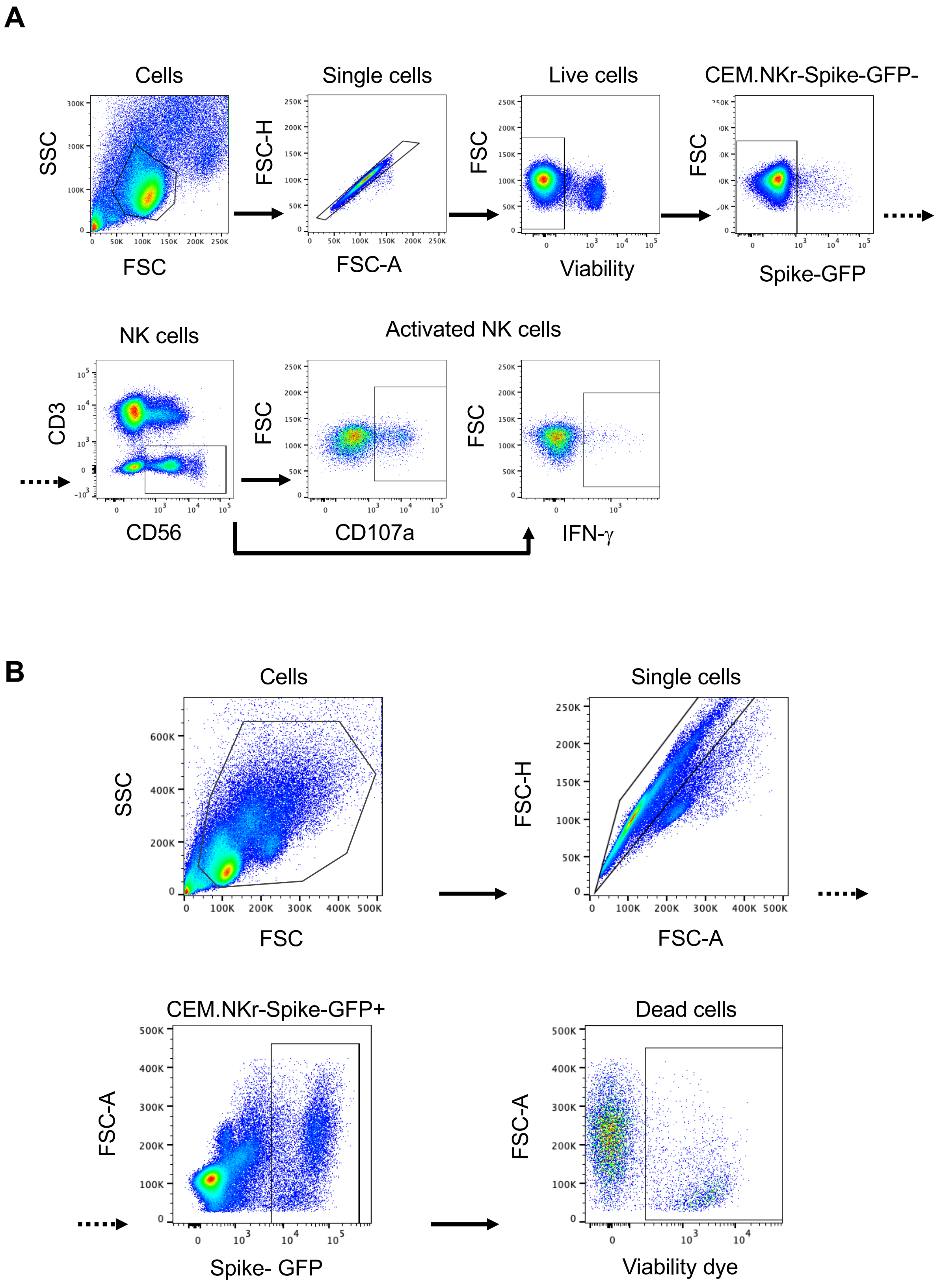
A
Cells
SSC
FSC
Single cells
FSC-H
FSC-A
Live cells
FSC
Viability
CEM.NKr-Spike-GFP-
FSC
Spike-GFP
NK cells
CD3
CD56
Activated NK cells
FSC
CD107a
FSC
IFN-γ
B
Cells
SSC
FSC
Single cells
FSC-H
FSC-A
CEM.NKr-Spike-GFP+
FSC-A
Spike- GFP
Dead cells
FSC-A
Viability dye

**S4 Fig.**

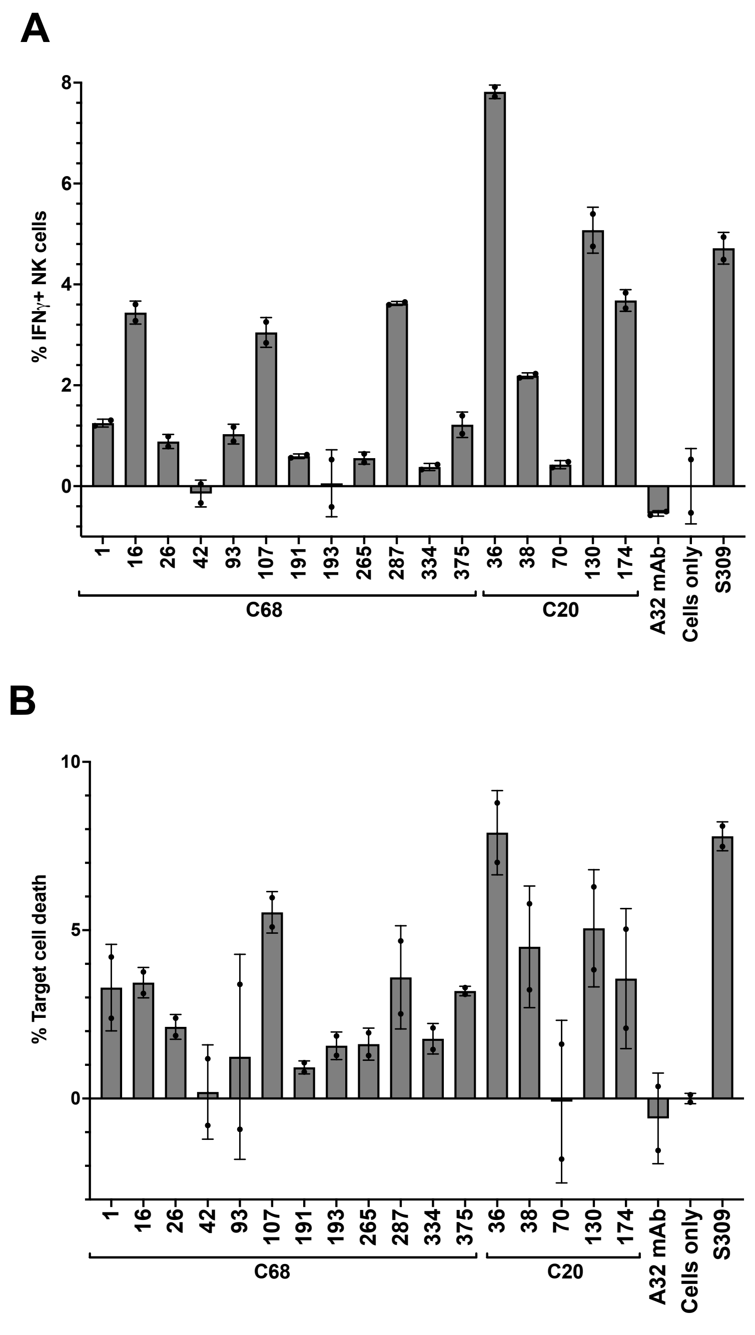
A
% IFNγ+ NK cells
8
6
4
2
0
1
16
26
42
93
107
191
193
265
287
334
375
36
38
70
130
174
A32 mAb
Cells only
S309
C68
C20
B
% Target cell death
10
5
0
1
16
26
42
93
107
191
193
265
287
334
375
36
38
70
130
174
A32 mAb
Cells only
S309
C68
C20

S5 Fig.

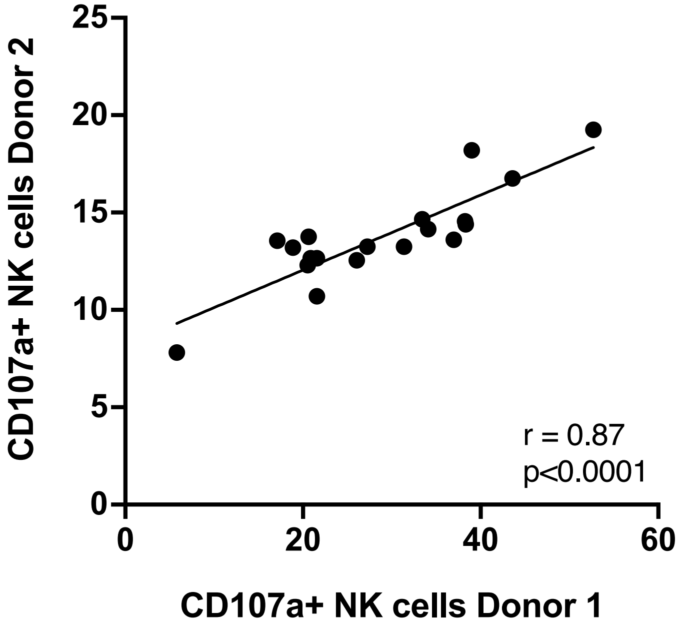

CD107a+ NK cells Donor 2
CD107a+ NK cells Donor 1
25
20
15
10
5
0
0
20
40
60
r = 0.87
p<0.0001


S6 Fig.

Biotinylated mAb (100ng/mL)

Blocking mAb (2500 ng/mL)

| | C68.1 | C68.14 | C68.16 | C68.23 | C68.26 | C68.35 | C68.40 | C68.42 | C68.43 | C68.49 | C68.81 | C68.93 | C68.107 | C68.109 | C68.144 | C68.191 | C68.193 | C68.204 | C68.228 | C68.251 | C68.265 | C68.287 | C68.334 | C68.337 | C68.375 | C20.38 | C20.59 | C20.70 | C20.130 | C13B8 | B6 | CV2-2164 | CV2-2333 |
|---|---|---|---|---|---|---|---|---|---|---|---|---|---|---|---|---|---|---|---|---|---|---|---|---|---|---|---|---|---|---|---|---|---|
| C68.1 | 98 | 0 | 0 | 2 | 0 | 101 | 0 | 89 | 0 | 0 | 0 | 8 | 0 | 0 | 100 | 0 | 0 | 22 | 0 | 94 | 3 | 99 | 6 | 0 | 100 | 3 | 0 | 101 | 19 | 0 | 0 | 97 | 0 |
| C68.4 | 0 | 0 | 0 | 5 | 0 | 0 | 44 | 0 | 0 | 0 | 0 | 0 | 0 | 0 | 0 | | 0 | 0 | 0 | 0 | 0 | 0 | 9 | 0 | 0 | 51 | 0 | | 0 | 0 | 0 | 26 | 0 |
| C68.5 | 2 | 0 | 0 | 1 | 0 | 1 | 17 | 0 | 13 | 0 | 2 | 0 | 0 | 0 | 12 | 14 | 0 | 0 | 6 | 0 | 1 | 0 | 5 | 23 | 4 | 12 | 0 | 26 | 0 | 0 | 0 | 8 | 49 |
| C68.14 | 0 | 90 | 10 | 0 | 0 | 86 | 4 | 0 | 0 | 11 | 0 | 87 | 3 | 0 | 95 | 0 | 0 | 0 | 13 | 85 | 6 | 0 | 2 | 0 | 0 | 21 | 0 | 100 | 0 | 0 | 5 | 92 | 0 |
| C68.16 | 2 | 0 | 100 | 1 | 0 | 1 | 1 | 1 | 25 | 5 | 1 | 7 | 80 | 0 | 17 | 0 | 3 | 0 | 40 | 2 | 7 | 0 | 2 | 0 | 0 | 53 | 0 | 14 | 38 | 0 | 12 | 0 | 0 |
| C68.23 | 0 | 76 | 0 | 60 | 98 | 2 | 0 | 0 | 0 | 100 | 16 | 0 | 59 | 94 | 0 | | 36 | 100 | 59 | 0 | 96 | 0 | 98 | 99 | 0 | 0 | 0 | | 52 | 0 | 0 | 0 | 0 |
| C68.26 | 0 | 12 | 29 | 53 | 100 | 0 | 0 | 0 | 0 | 100 | 0 | 12 | 39 | 97 | 5 | 0 | 0 | 100 | 20 | 9 | 89 | 0 | 100 | 100 | 4 | 28 | 0 | 12 | 36 | 0 | 8 | 0 | 0 |
| C68.35 | 54 | 95 | 22 | 1 | 3 | 95 | 43 | 86 | 0 | 0 | 0 | 95 | 27 | 0 | 99 | 13 | 0 | 14 | 10 | 98 | 0 | 2 | 0 | 0 | 96 | 25 | 0 | 100 | 18 | 0 | 1 | 100 | 0 |
| C68.40 | 4 | 9 | 4 | 3 | 16 | 4 | 87 | 1 | 0 | 0 | 14 | 0 | 0 | 0 | 0 | | 9 | 0 | 4 | 0 | 5 | 6 | 3 | 22 | 0 | 6 | 0 | | 6 | 0 | 0 | 1 | 12 |
| C68.42 | 98 | 87 | 0 | 0 | 0 | 101 | 2 | 96 | 5 | 0 | 9 | 99 | 0 | 0 | 100 | 0 | 0 | 69 | 0 | 99 | 0 | 100 | 0 | 0 | 99 | 5 | 0 | 101 | 24 | 0 | 2 | 99 | 2 |
| C68.43 | 0 | 0 | 1 | 0 | 0 | 0 | 0 | 0 | 65 | 1 | 30 | 19 | 0 | 0 | 8 | 73 | 0 | 13 | 1 | 0 | 3 | 1 | 0 | 0 | 22 | 20 | 0 | 32 | 34 | 0 | 6 | 0 | 75 |
| C68.49 | 0 | 0 | 25 | 6 | 58 | 2 | 0 | 1 | 0 | 97 | 9 | 11 | 24 | 41 | 3 | 0 | 0 | 97 | 19 | 0 | 47 | 0 | 81 | 91 | 0 | 31 | 0 | 14 | 40 | 0 | 14 | 0 | 0 |
| C68.72 | 2 | 11 | 0 | 0 | 0 | 4 | 0 | 1 | 0 | 53 | 0 | 0 | 0 | 0 | 0 | | 4 | 0 | 0 | 0 | 10 | 0 | 5 | 0 | 0 | 0 | 0 | | 0 | 0 | 0 | 0 | 0 |
| C68.81 | 2 | 14 | 0 | 1 | 0 | 2 | 38 | 1 | 9 | 2 | 77 | 0 | 0 | 0 | 0 | | 0 | 0 | 0 | 0 | 2 | 2 | 1 | 0 | 0 | 0 | 0 | | 99 | 0 | 0 | 24 | 0 |
| C68.93 | 0 | 99 | 28 | 3 | 0 | 99 | 14 | 91 | 0 | 4 | 10 | 99 | 37 | 0 | 99 | 0 | 0 | 10 | 35 | 97 | 10 | 0 | 3 | 26 | 96 | 28 | 0 | 101 | 24 | 0 | 21 | 97 | 12 |
| C68.97 | 50 | 91 | 7 | 0 | 0 | 92 | 12 | 71 | 50 | 0 | 0 | 81 | 6 | 0 | 98 | 24 | 4 | 32 | 0 | 93 | 2 | 7 | 0 | 0 | 82 | | 0 | 100 | 12 | 0 | 2 | 89 | 10 |
| C68.107 | 2 | 8 | 101 | 0 | 0 | 8 | 2 | 0 | 0 | 16 | 10 | 3 | 99 | 0 | 10 | 0 | 49 | 20 | 76 | 2 | 6 | 0 | 2 | 0 | 0 | 69 | 0 | 17 | 34 | 1 | 17 | 0 | 0 |
| C68.109 | 2 | 8 | 1 | 65 | 98 | 0 | 0 | 0 | 0 | 100 | 10 | 0 | 36 | 95 | 0 | 0 | 0 | 100 | 5 | 0 | 79 | 0 | 99 | 99 | 0 | 30 | 0 | 3 | 47 | 0 | 9 | 0 | 0 |
| C68.144 | 24 | 91 | 0 | 1 | 0 | 91 | 0 | 59 | 10 | 0 | 0 | 89 | 0 | 0 | 97 | 3 | 0 | 6 | 5 | 86 | 2 | 3 | 2 | 0 | 80 | 15 | 0 | 101 | 20 | 0 | 0 | 83 | 0 |
| C68.191 | 2 | 0 | 17 | 4 | 0 | 5 | 45 | 0 | 92 | 7 | 0 | 9 | 0 | 0 | 18 | 94 | 0 | 6 | 14 | 0 | 4 | 2 | 7 | 0 | 0 | 30 | 0 | 27 | 33 | 2 | 2 | 48 | 93 |
| C68.193 | 1 | 0 | 101 | 0 | 0 | 2 | 20 | 0 | 23 | 7 | 18 | 1 | 100 | 0 | 9 | 28 | 96 | 2 | 99 | 0 | 4 | 1 | 4 | 28 | 0 | 72 | 0 | 7 | 33 | 0 | 11 | 0 | 14 |
| C68.204 | 1 | 2 | 26 | 5 | 64 | 10 | 4 | 4 | 0 | 90 | 0 | 14 | 28 | 54 | 10 | 6 | 0 | 97 | 19 | 0 | 56 | 4 | 82 | 90 | 17 | 27 | 0 | 8 | 20 | 2 | 11 | 7 | 6 |
| C68.228 | 0 | 0 | 100 | 0 | 0 | 0 | 1 | 0 | 0 | 0 | 9 | 0 | 99 | 0 | 0 | | 88 | 0 | 92 | 0 | 0 | 0 | 0 | 0 | 0 | | 0 | | 39 | 0 | 0 | 0 | 0 |
| C68.251 | 90 | 95 | 0 | 1 | 2 | 99 | 0 | 91 | 8 | 0 | 0 | 97 | 11 | 0 | 99 | 10 | 0 | 65 | 0 | 98 | 0 | 16 | 9 | 0 | 99 | 14 | 0 | 101 | 7 | 0 | 0 | 97 | 12 |
| C68.265 | 0 | 0 | 62 | 92 | 101 | 0 | 0 | 0 | 0 | 100 | 0 | 2 | 0 | 99 | 4 | 0 | 0 | 100 | 2 | 0 | 97 | 0 | 101 | 100 | 0 | 11 | 7 | 0 | 35 | 0 | 8 | 0 | 0 |
| C68.287 | 80 | 0 | 11 | 3 | 0 | 24 | 41 | 0 | 0 | 0 | 0 | 10 | 0 | 0 | 84 | 9 | 0 | 95 | 11 | 3 | 7 | 85 | 7 | 0 | 95 | 28 | 0 | 101 | 23 | 0 | 10 | 95 | 0 |
| C68.334 | 2 | 0 | 9 | 55 | 91 | 0 | 15 | 0 | 0 | 99 | 9 | 0 | 27 | 94 | 8 | 0 | 0 | 99 | 28 | 0 | 72 | 1 | 99 | 99 | 3 | 30 | 0 | 7 | 19 | 0 | 11 | 2 | 8 |
| C68.337 | 3 | 6 | 0 | 54 | 99 | 2 | 6 | 3 | 0 | 100 | 5 | 0 | 59 | 96 | 0 | | 23 | 99 | 55 | 0 | 96 | 6 | 98 | 99 | 0 | 0 | 0 | | 37 | 0 | 7 | 0 | 2 |
| C68.375 | 75 | 15 | 3 | 0 | 3 | 95 | 0 | 89 | 0 | 5 | 5 | 93 | 25 | 0 | 97 | 5 | 0 | 97 | 20 | 91 | 6 | 84 | 3 | 0 | 97 | 27 | 0 | 101 | 16 | 0 | 6 | 93 | 0 |
| C20.36 | 5 | 18 | 0 | 0 | 9 | 2 | 0 | 9 | 0 | 0 | 5 | 2 | 0 | 0 | 0 | 0 | 1 | 0 | 0 | 0 | 10 | 1 | 0 | 0 | 0 | 1 | 0 | | 31 | 0 | 2 | 0 | 0 |
| C20.38 | 1 | 9 | 16 | 4 | 0 | 0 | 0 | 7 | 0 | 4 | 0 | 3 | 1 | 0 | 11 | 0 | 0 | 1 | 28 | 0 | 0 | 3 | 3 | 0 | 3 | 81 | 0 | 3 | 10 | 0 | 11 | 0 | 0 |
| C20.59 | 0 | 5 | 0 | 0 | 0 | 0 | 0 | 0 | 16 | 0 | 17 | 0 | 10 | 0 | 5 | 5 | 2 | 3 | 0 | 0 | 1 | 0 | 0 | 0 | 0 | | 52 | 2 | 9 | 1 | 3 | 1 | 0 |
| C20.70 | 6 | 35 | 0 | 4 | 0 | 4 | 40 | 10 | 0 | 22 | 0 | 22 | 0 | 0 | 0 | 0 | 0 | 0 | 0 | 0 | 4 | 1 | 0 | 0 | 2 | 0 | 0 | | 9 | 0 | 0 | 65 | 0 |
| C20.130 | 1 | 5 | 0 | 3 | 0 | 10 | 10 | 0 | 0 | 3 | 59 | 2 | 0 | 1 | 0 | | 0 | 0 | 0 | 0 | 3 | 0 | 0 | 41 | 0 | 0 | 0 | | 94 | 0 | 0 | 10 | 17 |
| C20.174 | 2 | 1 | 0 | 1 | 0 | 3 | 2 | 0 | 0 | 36 | 0 | 0 | 0 | 19 | 0 | | 0 | 0 | 0 | 0 | 5 | 0 | 16 | 77 | 0 | 0 | 0 | | 0 | 0 | 0 | 0 | 5 |
| B6 | 0 | 18 | 36 | 1 | 0 | 2 | 1 | 2 | 0 | 0 | 13 | 7 | 19 | 0 | 7 | 5 | 0 | 3 | 17 | 0 | 4 | 2 | 2 | 0 | 0 | 23 | 0 | 12 | 28 | 2 | 90 | 0 | 0 |
| CV2-2002 | 2 | 26 | 2 | 3 | 4 | 1 | 0 | 2 | 36 | 9 | 23 | 15 | 7 | 0 | 4 | 49 | 0 | 2 | 8 | 0 | 1 | 5 | 0 | 0 | 4 | 12 | 0 | 22 | 39 | 0 | 2 | 0 | 55 |
| CV2-2164 | 58 | 94 | 0 | 2 | 0 | 95 | 0 | 81 | 24 | 0 | 0 | 85 | 0 | 0 | 97 | | 0 | 42 | 0 | 83 | 0 | 4 | 0 | 0 | 82 | 53 | 0 | 0 | 9 | 0 | 0 | 85 | 0 |
| CV2-2333 | 1 | 7 | 0 | 2 | 25 | 3 | 37 | 2 | 80 | 5 | 0 | 5 | 57 | 7 | 0 | | 26 | 0 | 53 | 0 | 21 | 2 | 1 | 0 | 0 | 0 | 0 | 0 | 47 | 0 | 3 | 50 | 90 |
| CV3-25 | 1 | 12 | 13 | 0 | 6 | 0 | 16 | 0 | 10 | 8 | 18 | 17 | 8 | 4 | 0 | 0 | 0 | 4 | 5 | 0 | 0 | 0 | 1 | 30 | 6 | 29 | 0 | 4 | 25 | 0 | 71 | 10 | 18 |
| 76E1 | 1 | 4 | 0 | 0 | 0 | 0 | 0 | 0 | 12 | 12 | 0 | 0 | 10 | 0 | 6 | 0 | 0 | 1 | 0 | 1 | 0 | 0 | 2 | 0 | 9 | | 0 | 0 | 12 | 9 | 3 | 0 | 1 |
| VP12E7 | 0 | 4 | 0 | 0 | 0 | 0 | 3 | 0 | 8 | 8 | 4 | 4 | 8 | 1 | 1 | 7 | 2 | 2 | 0 | 2 | 2 | 2 | 0 | 1 | 2 | | 0 | 0 | 5 | 6 | 2 | 0 | 4 |
| C13B8 | 1 | 28 | 4 | 0 | 0 | 1 | 0 | 7 | 33 | 33 | 30 | 30 | 12 | 0 | 2 | 12 | 5 | 7 | 0 | 9 | 3 | 3 | 4 | 3 | 0 | | 0 | 0 | 40 | 83 | 2 | 0 | 4 |

### S7 Fig.

| Comp Group | mAb ID | IGHV | IGLV or IGKV | Heavy %SHM | Heavy CDR3 length |
|---|---|---|---|---|---|
| Group 1 | **C68.1** | HV5-10-1 | KV4-1 | 5.0 | 13 |
| | **C68.14** | HV1-69 | KV2-30 | 4.0 | 20 |
| | **C68.35** | HV1-69 | KV3-11 | 1.4 | 15 |
| | **C68.42** | HV3-30 | KV1-5 | 4.0 | 18 |
| | **C68.93** | HV4-31 | KV3-20 | 3.2 | 16 |
| | **C68.97** | HV1-69 | KV3-11 | 1.4 | 15 |
| | **C68.144** | HV1-69 | KV3-11 | 2.2 | 15 |
| | **C68.251** | HV1-69 | KV3-11 | 3.0 | 15 |
| | **C68.287** | HV1-46 | KV1-5 | 6.3 | 14 |
| | **C68.375** | HV3-30 | KV3-20 | 4.6 | 9 |
| | **C20.70** | HV3-23 | LV2-14 | 2.5 | 14 |
| Group 2 | **C68.23** | HV1-2 | KV4-1 | 2.5 | 15 |
| | **C68.26** | HV3-30 | LV1-47 | 3.6 | 13 |
| | **C68.49** | HV3-30 | KV1-5 | 3.6 | 14 |
| | **C68.109** | HV3-30 | KV3-20 | 3.6 | 14 |
| | **C68.204** | HV3-21 | KV1-NL1 | 4.6 | 17 |
| | **C68.265** | HV3-30 | KV1D-39 | 3.6 | 14 |
| | **C68.334** | HV4-59 | KV2D-28 | 4.7 | 14 |
| | **C68.337** | HV4-34 | KV1D-39 | 5.9 | 23 |
| | **C20.174** | HV3-30-3 | LV1-40 | 3.7 | 11 |
| Group 3 | **C68.16** | HV1-46 | KV3-20 | 2.8 | 13 |
| | **C68.107** | HV3-7 | KV1-12 | 4.1 | 22 |
| | **C68.193** | HV3-7 | LV3-19 | 1.5 | 24 |
| | **C68.228** | HV3-7 | KV3-15 | 3.4 | 22 |
| | **C20.38** | HV3-23 | LV8-61 | 2.8 | 24 |
| Group 4 | **C68.5** | HV1-3 | LV3-10 | 9.7 | 13 |
| | **C68.40** | HV3-7 | LV3-21 | 2.5 | 13 |
| | **C68.43** | HV3-74 | LV3-1 | 3.0 | 14 |
| | **C68.191** | HV3-23 | LV3-10 | 3.7 | 10 |
| Group 5 | **C20.36** | HV3-53 | KV1-5 | 0.3 | 9 |
| | **C20.59** | HV3-30-3 | LV3-1 | 1.6 | 15 |
| | **C20.130** | HV3-30-3 | LV3-10 | 4.9 | 14 |
| | **C68.4** | HV4-59 | KV4-1 | 0.5 | 21 |
| | **C68.72** | HV1-2 | LV1-47 | 1.3 | 21 |
| | **C68.81** | HV3-30 | LV3-10 | 4.1 | 15 |

**S8 Fig**.

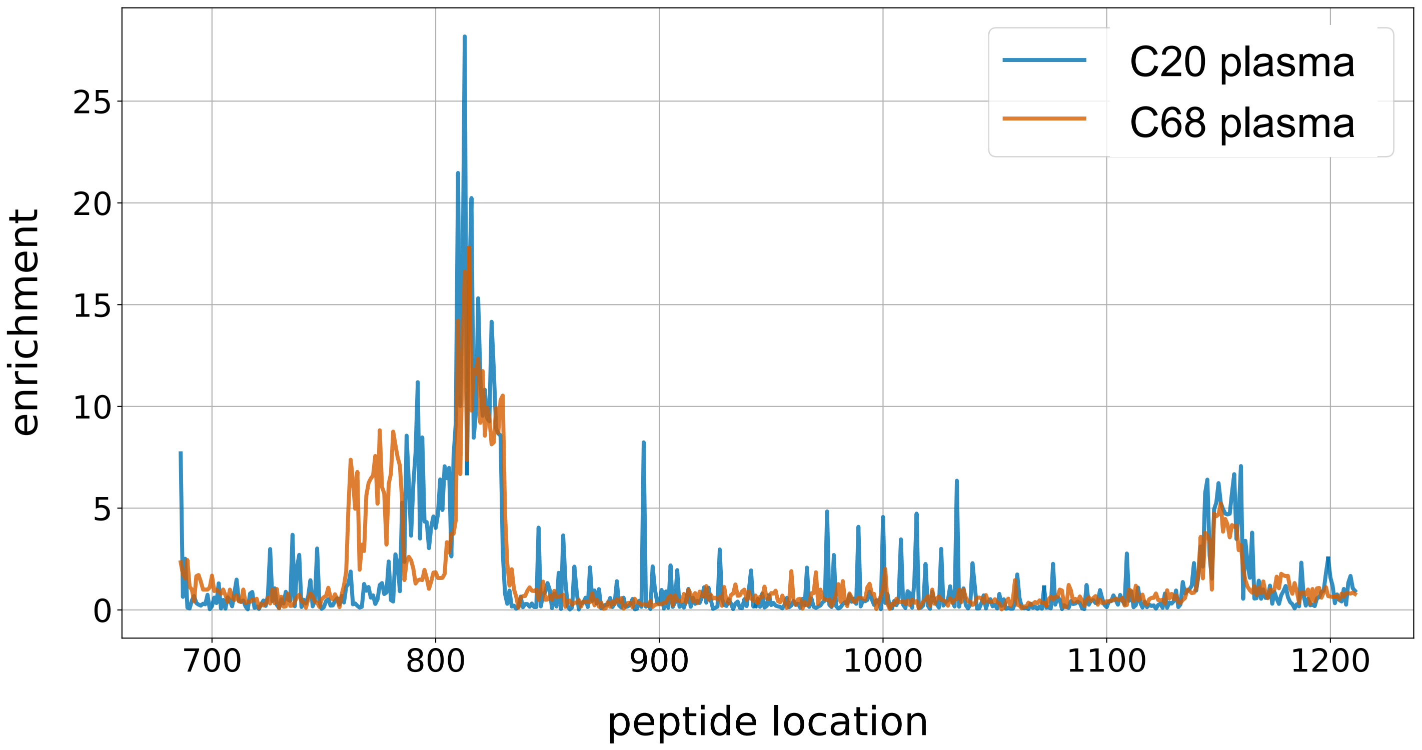

C20 plasma
C68 plasma
enrichment
0
5
10
15
20
25
700
800
900
1000
1100
1200
peptide location


**S9 Fig.**

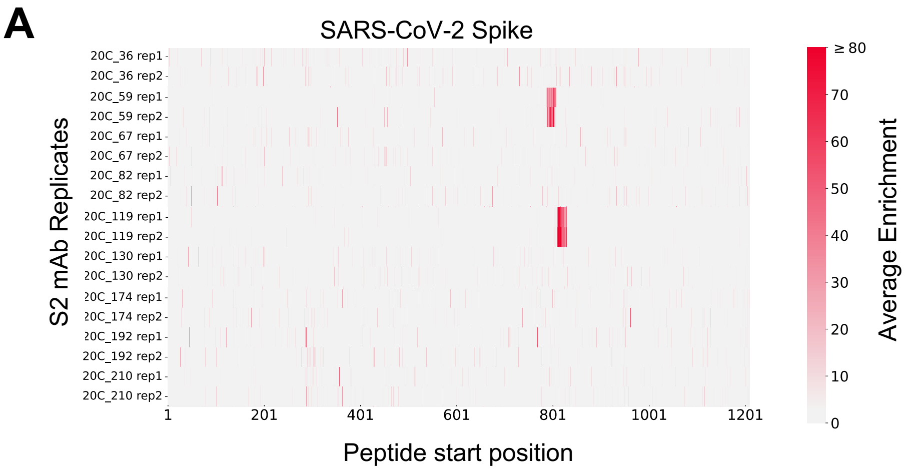
A
SARS-CoV-2 Spike
S2 mAb Replicates
20C_36 rep1
20C_36 rep2
20C_59 rep1
20C_59 rep2
20C_67 rep1
20C_67 rep2
20C_82 rep1
20C_82 rep2
20C_119 rep1
20C_119 rep2
20C_130 rep1
20C_130 rep2
20C_174 rep1
20C_174 rep2
20C_192 rep1
20C_192 rep2
20C_210 rep1
20C_210 rep2
1
201
401
601
801
1001
1201
Peptide start position
≥ 80
70
60
50
40
30
20
10
0
Average Enrichment

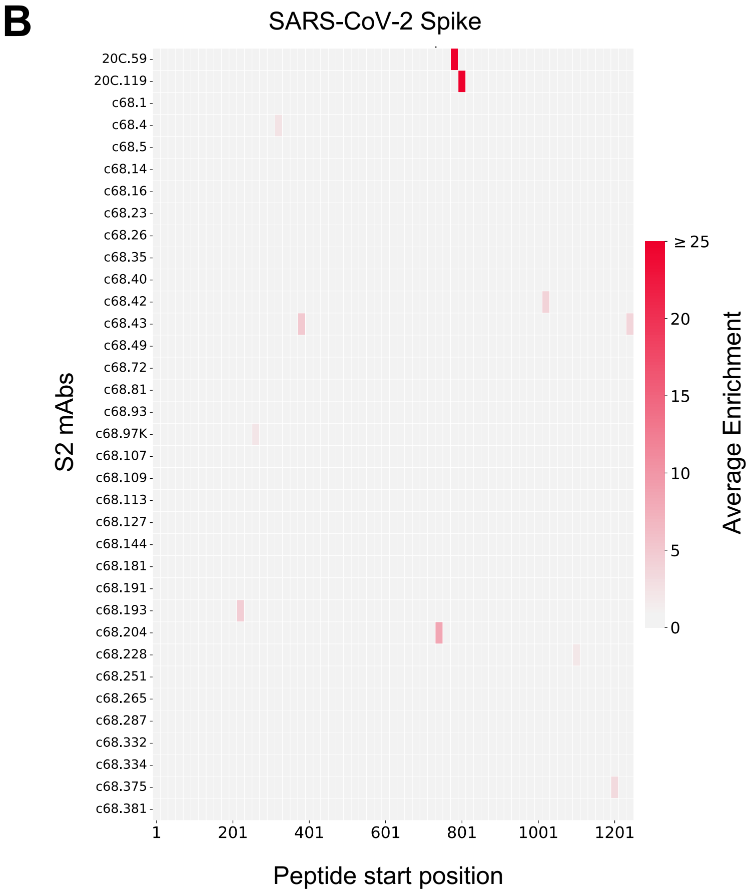
B
SARS-CoV-2 Spike
S2 mAbs
20C.59
20C.119
c68.1
c68.4
c68.5
c68.14
c68.16
c68.23
c68.26
c68.35
c68.40
c68.42
c68.43
c68.49
c68.72
c68.81
c68.93
c68.97K
c68.107
c68.109
c68.113
c68.127
c68.144
c68.181
c68.191
c68.193
c68.204
c68.228
c68.251
c68.265
c68.287
c68.332
c68.334
c68.375
c68.381
1
201
401
601
801
1001
1201
Peptide start position
≥ 25
20
15
10
5
0
Average Enrichment

**S10 Fig.**

**A**

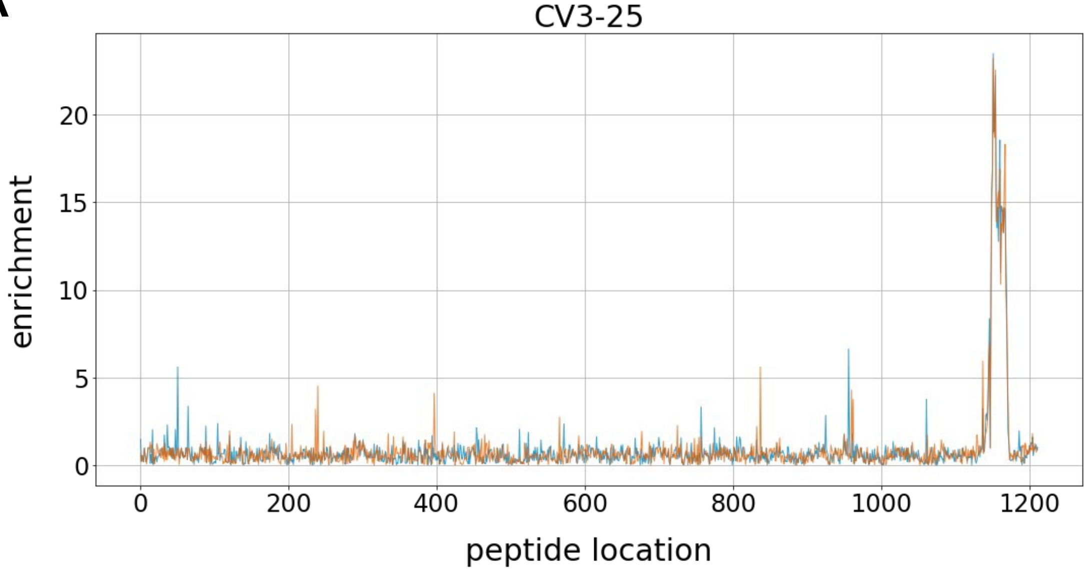
CV3-25
enrichment
0
5
10
15
20
0
200
400
600
800
1000
1200
peptide location

**B**

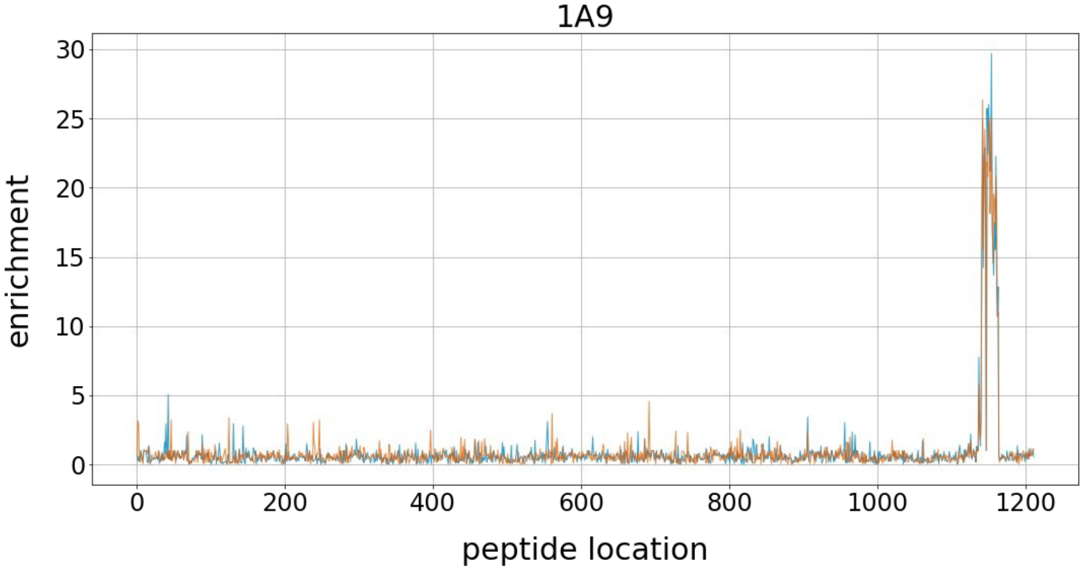
1A9
enrichment
0
5
10
15
20
25
30
0
200
400
600
800
1000
1200
peptide location

**S11 Fig.**

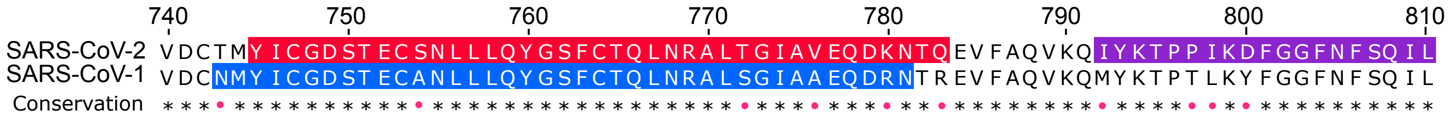
740
750
760
770
780
790
800
810
SARS-CoV-2 VDCTMYICGDSTECSNLLLQYGSFCTQLNRALTGIAVEQDKNTQEVFAQVKQIYKTPPIKDFGGFNFSQIL
SARS-CoV-1 VDCNMYICGDSTECANLLLQYGSFCTQLNRALSGIAAEQDRNTREVFAQVKQMYKTPTLKYFGGFNFSQIL
Conservation

**S12 Fig.**

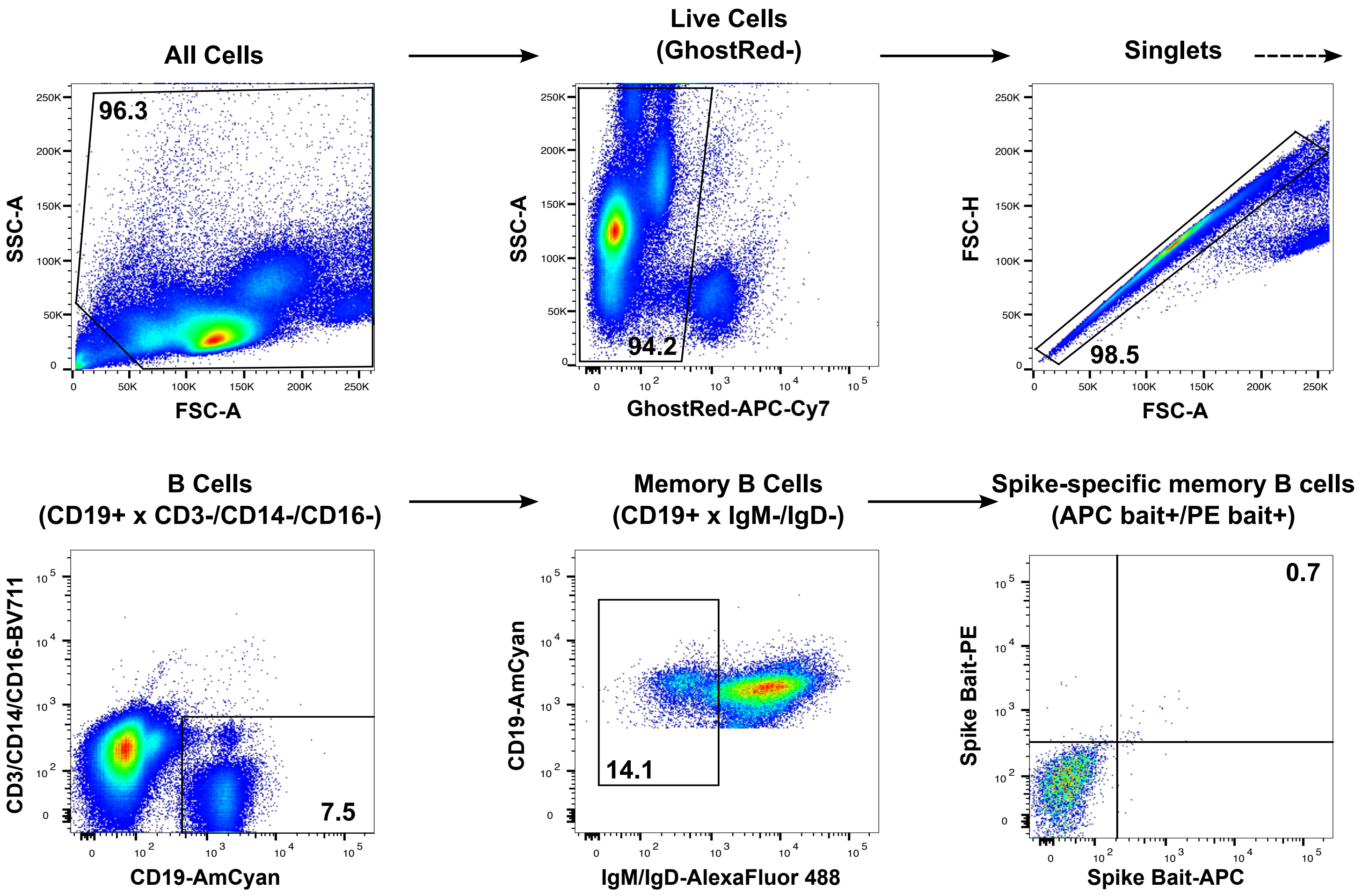
All Cells
Live Cells (GhostRed-)
Singlets
96.3
94.2
98.5
SSC-A
FSC-A
GhostRed-APC-Cy7
FSC-H
B Cells (CD19+ x CD3-/CD14-/CD16-)
Memory B Cells (CD19+ x IgM-/IgD-)
Spike-specific memory B cells (APC bait+/PE bait+)
7.5
14.1
0.7
CD3/CD14/CD16-BV711
CD19-AmCyan
IgM/IgD-AlexaFluor 488
Spike Bait-PE
Spike Bait-APC

S13 Fig.
