## Supplemental Table for "The S2 subunit of spike encodes diverse targets for functional antibody responses to SARS-CoV-2"

| | mAb ID | IGHV gene | IGHD gene | IGHJ gene | CDR3 length (aa) | %SHM | IGKV or IGLV gene | IGKJ or IGLJ gene | CDR3 length (aa) | %SHM | IGH variable region aa seq | IGK or IGL variable region aa seq |
|---|---|---|---|---|---|---|---|---|---|---|---|---|
| C68 | 1 | IGHV5-10-1*01 | IGHD6-6*01 | IGHJ4*02 | 13 | 5.0 | IGKV4-1*01 | IGKJ3*01 | 9 | 3.2 | EVQLVQSGAGVRKPGESLRISCKGSGYNFTSYWITWVRQMPGKGLEWMGRIDPTDSYTTYSPAFQGHVTISADKSISTAYLQWSRLKASDTAMYYCARHVLKSAALLGYWGQGTLVTVSS | DIVMTQSPDSLAVSLGERATINCKSSQSVLYSSSNKNYLGWYQQKPGQPPKLLIYWASTRESGVPDRFTGSGSGTDFTLTISSLQPEDVAVYYCQQYFLPPFTFGPGTTVNIK |
| | 4 | IGHV4-59*01 | IGHD6-19*01 | IGHJ6*03 | 21 | 0.5 | IGKV4-1*01 | IGKJ2*01 | 11 | 1.7 | QVQLQESGPGLVKPSETLSLTCTVSGGSISSYYWSWIRQPPGKGLEWIGYIYYSGSTNYNPSLKSRVTISVDTSKNQFSLKLSSVTAADTAVYYCARERKTEYSSGWYYYYYMDVWGKGTTVTVSS | DIVMTQSPDSLAVSLGERATINCKSSQSLSYTSSKKNYLAWYQQKAGQPPKLLIYWASTRESGVPDRFSGSGSGTDFTLTISSLQAEDVAVYYCQQYYSXPPVYTFGQGTKLEIK |
| | 5 | IGHV1-3*01 | IGHD6-13*01 | IGHJ4*02 | 13 | 9.7 | IGLV3-10*01 | IGLJ2*01 | 10 | 3.3 | QVQLVQSGAEVKKPGASVKVSCKASGNSFTSYTFQWVRQAPGQRLEWMGWVNGGNGNTKYSQKFQGRVTIFRDTSVSTVYMRLNSLRYEDTGVYYCASGEMAAAGLFDFWGQGTLVTVSS | SYELTQPPSVSVSPGQTARITCSGDALPKKYTYWYQQKSGQAPVLVIYEDNTRPSGIPERFSGSSSGTMATLTISGAQVEDEADYYCYSTDYSGQGVFGG |
| | 14 | IGHV1-69*01 | IGHD6-19*01 | IGHJ4*02 | 20 | 4.0 | IGKV2-30*02 | IGKJ4*01 | 9 | 0.3 | QVQLVQSGAEVKKPGSSVKVSCKVSGGTFSSYSISWVRQAPGQGLEWMGGIIPIFGIVNYAQKFQGRVTVTADKSTSTAYMELSSLRSEDTAVYYCARGLSSGWFAVAAQPPFFDSWGQGTLVTVSS | DVVMTQSPLSLPVTLGQPASXSCRXSQSLVHSDGNTYLNWFQQRPGQSPRRLIYKVSNRDSGVPDRFSGSGSGTDFTLKISRVEAEDVGVYYCMQGTHWPLTFGGGTKVEIK |
| | 16 | IGHV1-46*01 | IGHD1-26*01 | IGHJ4*02 | 13 | 2.8 | IGKV3-20*01 | IGKJ1*01 | 9 | 1.5 | QVQLVQSGAEVKKPGASVKVSCKASGYTFTSYYIHWVRQAPGQGLEWMGVIIPSGGSTTYAQRFQGRVTMSRDTSTSTVYMELSSLRSEDTAVYYCAGTGIVVNWVFDYWGQGTLVTVSS | EIVLTQSPGTLSLSPGERATLSCRASQSVSSYLAWYQQKPGQAPRLLIYGASSRATGIPDRVSGSGSGTDFTLTISRLEPEDFAVYYCQQYRSSPPTFGQGTKVEIK |
| | 23 | IGHV1-2*02 | IGHD3-10*01 | IGHJ4*02 | 15 | 2.5 | IGKV4-1*01 | IGKJ2*01 | 11 | 1.7 | QVQLVQSGAEVKKPGASVKVSCKASGYTFIGYFMHWVRQTPGQGLEWMGWINPNSGTTNYAQKFQGRVTMTRDTSISTVYMELSRLRSDDTAVYYCARGQRWSGELIQFDYWGQGTLVTVSS | DIVMTQSPDSLAVSLGERATIKCKSSQSVLYSSSNKNYLAWYQQKPGQPPKLLIYWASTRESGVPDRFSGSGSGTDFTLTISSLQAEDVAVYYCQQYYSTPLMYTFGQGTKLEIK |
| | 26 | IGHV3-30*18 | IGHD5-18*02 | IGHJ4*02 | 13 | 3.6 | IGLV1-47*01 | IGLJ1*01 | 12 | 3.3 | EVQLVESGGGVVQPGRSLRLSCAASGFTFTTYAMHWFRQAPGKGLEWVALISYDGINKYYPDSVRGRFTISRDNSKNTLFLQMNSLRVEDTAVYYCAKVAYSGYDLPDYWGQGTLVTVSS | MPVLTQPPSASGTPGQRVTISCSGSSSNIGSNYVYWYQHLPGTAPKLLIYRNDQRPSGVPDRFSGSKSGTSASLAISGLRSEDEADYYCATWGVSVSGNYVFGAGTKVTVL |
| | 35 | IGHV1-69*01 | IGHD4-17*01 | IGHJ4*02 | 15 | 1.4 | IGKV3-11*01 | IGKJ5*01 | 11 | 2.1 | QVQVVQSGAEVKKPGSSVKVSCKASGGTFSSYAISWVRQAPGQGLEWMGGVIPIFGTTNYAQKFQGRVTITADESTSTAYMELSSLRSEDTAVYYCARVLGWDGDSGGTTNWGQGTLVTVSS | EIVLTQSPATLSLSPGERATLSCRASQSVSNFLAWFQHKPGQAPRLLIYDASNRATGIPARFSGSGSGTDFTLTISSLEPEDFAVYYCQHRRNWPPVITFGQGTRLEIK |
| | 40 | IGHV3-7*03 | IGHD3-9*01 | IGHJ4*02 | 13 | 2.5 | IGLV3-21*03 | IGLJ3*02 | 12 | 3.4 | EVQLVESGGGLVQPGGSLRLSCAVSGVIFSSYWMSWVRQAPGKGLEWVANINQDGSEKYYVDSVKGRFTISRDNAKNSLYLQMNSLRADDTAVYYCARGGYFDWSVSDYWGQGTLVTVSS | QPVLTQPPSVSVAPGKTARITCGGDNIGSKSVHWYQQKPGQAPVLVVYDDSDRPSGIPERFSGSKSGNTATLTISRVEAGDEADYYCQVWDNSSDLLWVFGGGTKLTVL |
| | 42 | IGHV3-30*18 | IGHD6-19*01 | IGHJ4*02 | 18 | 4.0 | IGKV1-5*01 | IGKJ2*01 | 9 | 1.9 | EVQLVESGGGVVQPGRSLRLSCAAFGFTFSSYGIHWVRQTPGKGLEWVAVISNDGSKKYYADSVKGRFTLSRDNSKNTLYLQMNSLRTEDAAIYYCAKVENIVAAGPTWGYFDYWGQGALVTVSS | DIQLTQSPSTLSASVGDRVTITCRASQTISSWLAWYQQKPGKAPKLLIYDASNLESGVPSRFSGSGSGTEFTLTISSLQPDDFATYYCQQYNRYPYTFGQGTKLEIK |
| | 43 | IGHV3-74*01 | IGHD3-3*01 | IGHJ6*03 | 14 | 3.0 | IGLV3-1*01 | IGLJ2*01 | 9 | 5.0 | EVQLVESGGGLVQPGGSLRLSCAASGFTFSSYWMHWVRQAPGKGLVWVSRITSDAYSTSYADSVKGRFTISRDNAKNTLHLQMNSLRAGDTAVYYCVRATVMASSYHMDVWGKGTTVTVSS | MPVLTQPPSVSVSPGQPASITCSGEKLGDKFVCWYQQKPGQSPVLVIYQDTKRPSGIPERFSGSNSANTATLTISGTQAMDEADYYCQAWDSSTGVFGGGTKLTVL |
| | 49 | IGHV3-30*18 | IGHD3-10*01 | IGHJ6*02 | 14 | 3.6 | IGKV1-5*01 | IGKJ1*01 | 8 | 1.6 | QVQLVQSGGGVVQPGRSLRLSCAASGFTFSIYGMHWVRQGPGKGLEWVALISYDGSNNYYTDSVKGRFTISRDNSKNTLYLQMNSLRLEDTAVYYCARPRSGSYFNGMDVWGQGTTVTVSS | DIQMTQSPSTLSASVGDRVTITCRASQSINSRLAWYQQKPGKAPKLLIYDASSLESGVPSRFRGSGSGTEFTLAISSLQPDDFATYYCQQYNTYWTFGQGTKVEIK |
| | 72 | IGHV1-2*02 | IGHD3-10*01 | IGHJ6*02 | 21 | 1.3 | IGLV1-47*01 | IGLJ3*02 | 11 | 1.9 | QVQLVQSGAEMKKPGASVKVSCKASGYTFTGYYMHWVRQAPGQGLEWMGWINPNSGGTNYAQNFQGRVTMTRDTSISTAYMELSSLRSDDTAVYFCASYGSGSYYNGDYYYYYGMDVWGQGTTVTVSS | QPVLTQPPSASGTPGQTVTISCSGSSSNIGSNYVYWYQQLPGTAPKLLIYGNNQRPSGVPDRFSGSKSGTSASLAISGLRSEDEADYYCAAWDDTLSGWVFGGGTK |
| | 81 | IGHV3-30*18 | IGHD6-19*01 | IGHJ6*02 | 15 | 4.1 | IGLV3-10*01 | IGLJ3*02 | 11 | 1.2 | EVQLVESGGGVVQPGRSLRLSCAASGFNFNNYGMHWVRQAPGKGLEWVAVIWYDGTNKYYADSVKGRFTISRDNSKNTLYLQLNSLRAEDTAVYYCARDFGFSSAFDFADVWGQGTTVTVSS | SYELTQPPSVSVSPGQTARITCSGDALPKKYAYWYQQKSVQAPLLVIYEDSKRPSGIPERFSGSSSGTMATLTISGAQVEDEADYYCYSTDSSGNHWVFGGGTKLTVL |
| | 93 | IGHV4-31*03 | IGHD2-21*02 | IGHJ5*02 | 16 | 3.2 | IGKV3-20*01 | IGKJ2*01 | 9 | 1.9 | QVQLQESGPRLVKPSQTLSLICTVSGGSISSDGYYWSWVRQHPGRGLEWIGYIFYTGSTYYNPSLKSRVTISVDTSKNQFSLNLNSVAAADTAVYYCARGVFCGADCYNWFDPWGQGTLVTVSS | EIVLTQSPGTLSLSPGERATLSCRASQSVSSSYIAWYQQKPGQAPRLLIYGASSRATGIPDRFSGSGSGTDFTLTISRLEPEDFAVYYCQQFGTSRTTFGQGTKLEIK |
| | 97 | IGHV1-69*01 | IGHD3-22*01 | IGHJ4*02 | 15 | 1.4 | IGKV3-11*01 | IGKJ2*01 | 11 | 0.9 | QVQLVQSGAEVKKPGSSVKVSCKASGGTFSSYAISWVRQAPGQGPEWMGGIIPIFGTTNYAQKFQGRVTITADESTSTAYMELSSLRSEDTAVYYCVRIDQYDSSGYYLDYWGQGTLVTVSS | EIVLTQSPATLSLSPGERATLSCRASQSVSSFLAWYQQKPGQAPRLLIYDASNRATGIPARFSGSGSGTDFTLTISSLEPEDFAVYYCHQRSNWPPRYTFGQGTRLEIK |
| | 107 | IGHV3-7*03 | IGHD6-6*01 | IGHJ6*03 | 22 | 4.1 | IGKV1-12*01 | IGKJ3*01 | 10 | 3.7 | EVQLVESGGGLVQPGGSLRLSCVASGFTFSSYWMSWVRQVPGQGLEWVANIKQDGSEKYYVDSVKGRFTISRDNAKNSLFLQMNSLRAEDTAVYYCARKNIAALLQNRGYYYYYYMDVWGKGTTVTVSS | DIQLTQSPSSVSASVGDRVAITCRASQDISSRLAWYQQKPGKAPKLLIYGASSLQSGVPSRFSGSGSGTDFTLTISSLQPEDFATYYCQQAHSFLRSTFGPGTKVDIK |
| | 109 | IGHV3-30*18 | IGHD1-26*01 | IGHJ3*02 | 14 | 3.6 | IGKV3-20*01 | IGKJ2*01 | 10 | 2.7 | EVQLVESGGGVVQPGRSLRLSCAASGFIFSTYGMHWVRQAPGKGPEWVALISYDGYNKYYADSVRGRFTISRDNSKNTLYLQMNSLRVEDTAVYYCASKFGGSYLGAFDIWGQGTMVTVSS | EIVLTQSPGTLSLSPGERATLSCRASQSVSKNYLAWYQQKPGQAPRLLISGASSRATGIPDRFSGSGSGTDFTLTITRLEPEDFAVYYCQQYGSSPMYTFGQGTKLEIK |
| | 144 | IGHV1-69*01 | IGHD5-18*02 | IGHJ4*02 | 15 | 2.2 | IGKV3-11*01 | IGKJ4*01 | 11 | 1.2 | QVQLVQSGAEVKKPGSSVKVSCKASGGIFSSYAITWVRQAPGQGLEWMGGIIPIFGTANYAQKFQGRVTITADESTSTAYVDLSSLRSEDTAVYYCVRISAYDSGGSYFDYWGQGTLVTVSS | QVQLVQSGAEVKKPGSSVKVSCKASGGIFSSYAITWVRQAPGQGLEWMGGIIPIFGTANYAQKFQGRVTITADESTSTAYVDLSSLRSEDTAVYYCVRISAYDSGGSYFDYWGQGTLVTVSS |
| | 191 | IGHV3-23*01 | IGHD3-10*01 | IGHJ4*02 | 10 | 3.7 | IGLV3-10*01 | IGLJ2*01 | 5 | 1.0 | EVQLLESGGGLVQRGGSLRLSCAASGFTFSNYAMSWVRQAPGKGLEWVSAISDTGGSTYYADSVKGRFTIFRDNSKNTLYLQMNSLRAEDTAVYYCANQWFGGVEYWGQGTLVTVSS | SYELTQPPSVSVSPGQTARITCSGDALPKKYAYWYQQKSGQAPVLVIYEDSRRPSGIPERFSGSSSGTMATLTISGAQVEDEGDYYCYSPRVFGGGTKLTVL |
| | 193 | IGHV3-7*03 | IGHD2-2*01 | IGHJ6*03 | 24 | 1.5 | IGLV3-19*01 | IGLJ3*02 | 13 | 0.6 | EVQLVESGGGLVQPGGSLRLSCAASGFTFSSYWMNWVRQAPGKGLEWVANIKQDGREKYYVDSVKGRFTISRDNAKNSLYLQMNSLRAEDTAVYFCARFVVVPAAVGGGWLGYYYYYMDVWGKGTTVTVS | SSELTQDPAVSVALGQTVRITCQGDSLRSYYANWYQQKPGQAPALVIYGKNNRPSGIPDRFSGSSSGNTASLTITGAQAEDEADYYCNSRDSSGNHLDWVFGGGTKLTVL |
| | 204 | IGHV3-21*01 | IGHD6-13*01 | IGHJ6*03 | 17 | 4.6 | IGKV1-NL1*01 | IGKJ2*01 | 9 | 0.9 | EVQLVESGGGLVKPGGSLRLSCAASGFSFSTYTLNWVRQAPGKGLEWVSSISSTSSYIYYADSVKGRFTISRDNVKKSLYLQMNSLRAEDTAVYFCARGISSSWTPRGFYMDVWGNGTTVTVSS | DIQLTQSPSSLSASVGDRVTITCRASQGISNSLAWYQQRPGKAPKLLLYAASRLESGVPSRFSGSGSGTDYTLTISSLQPEDFATYYCQQFYSTPYTFGQGTKLEIK |
| | 228 | IGHV3-7*03 | IGHD3-10*01 | IGHJ6*02 | 22 | 3.4 | IGKV3-15*01 | IGKJ2*01 | 10 | 2.5 | EVQVVESGGGLVQPGGSLRLSCAASGFTFSSYWMSWVRQAPGKGLEWVANIRHDGREKYYVDSVKGRFTISRDKAKNSLYLQMNSLRAEDTAVYYCAREVRGLVDLEPPYSYYYAFDVWGQGTTVTVSS | EIVLTQSPATLSMSPGERATLSCRASQSVSSNLAWYQHKPGQAPRLLIYGASTRASGFPARFSGSGSGTEFTLTISSLQSEDFAVYYCQQYNNWPQYTFGQGTKLEIK |
| | 251 | IGHV1-69*01 | IGHD3-22*01 | IGHJ4*02 | 15 | 3.0 | IGKV3-11*01 | IGKJ5*01 | 11 | 2.4 | QVQVVQSGAEVKKPGSSVKVSCKTSGGTFSSYAISWVRQAPGQGLEWMGGIIPIFGTTRYAQKFQGRVTITADESTSTAYMELSSLRSEDTAMYYCANVGPYDSSGYYVDHWGQGTLVTVSS | EIVLTQSPPTLSLSPGERATLSCRASQSVSSFLAWYQQIPGQAPRLLIYDASNRATGIPARFSGSGSGTDFTLTISSLEPEDFAVYYCLHRSNWPPEITFGQGTRLEIK |
| | 265 | IGHV3-30*18 | IGHD6-13*01 | IGHJ6*02 | 14 | 3.6 | IGKV1D-39*01 | IGKJ3*01 | 9 | 2.5 | EVQLVESGGGVVQPGRSPRLSCAASGFTFSSNAMHWVRQAPGKGLEWVALISYDGDNKYYADSVKGRFIISRDNSKNTLYLQMSSLRAEDTAVYYCAKATAGSYYYGMDVWGQGTTVTVSS | DIQLTQSPSSLSASVGDRVTITCRASQSISNYLNWYQQKSGKAPKLLIYAASSLQSGVPSRFSGSGSGTDFTLTISSLQPEDFATYYCQQGFPTPFTFGPGTKVDIK |
| | 287 | IGHV1-46*01 | IGHD6-19*01 | IGHJ4*02 | 14 | 6.3 | IGKV1-5*01 | IGKJ4*01 | 10 | 4.6 | QVQLVQSGAEVKKPGASVKVSCKASGYTFTKYYIHWVRQAPGQGLEWMGIINPSGAGTIYAQNFQGRVTMTVDTSTSTVYMEVSSLKSEDTAVYYCARGQEQFLEQFFDYWGQGTQVTVSS | DIQLTQSPSTLSASVGDRVTITCRASQNIGNWLAWYQQKPGKAPNLLIYDASSLQSGIPSRFSGTGSGTDFTLTISSLQPDDFATYYCQQFKNYSPLTFGGGTKVEIK |
| | 334 | IGHV4-59*01 | IGHD3-10*01 | IGHJ6*03 | 14 | 4.7 | IGKV2D-28*01 | IGKJ1*01 | 9 | 1.2 | QVQLQESGPGLVKPSETLSLTCTVSGGSINNYFWSWIRQPPGKGLEWIGHMYYSGSTEYIPSLKSRVTMSVDTSKNQCSLKLSSVTAADTAVYFCARVLYGSGSYSMDVWGKGTTVTVSS | DIVMTQSPLSLPVTPGEPASISCRSSQSLLHSDGYNYLDWYLQKPGQSPQLLIYLGSNRASGVPDRFSGSGSGTDFTLKISRVEAEDVGVYYCMQSLQTPLTFGQGTRVEIK |
| | 337 | IGHV4-34*01 | IGHD6-13*01 | IGHJ5*02 | 23 | 5.9 | IGKV1D-39*01 | IGKJ3*01 | 9 | 5.9 | QVQLQQWGTGLLKSSETLSLTCAVYGESFSGYYWSWIRQSPGKGLEWVGEINHSGRTNYNPSLKSRVTISIDTSKNQFSLKLNSVTAADTAVYYCARNCHHFGISASGTSGFIKRFDSWGQGTLVTVSS | DIQMTQSPSPLSASIGDRVTITCRASQDINIYLNWYQQKPGKAPNLLIYGASTLQSGVPSRFSGSGSGTDFTLTISSLQPEDFATYYCQQSYSTLLTFGPGTTVDSK |
| | 375 | IGHV3-30*18 | IGHD3-10*01 | IGHJ4*02 | 9 | 4.6 | IGKV3-20*01 | IGKJ1*01 | 9 | 1.2 | EVQLVESGGGVVQPGRSLRLSCAASGFTFNNYGMHWVRQAPGKGLDWVAAIWHDGSDKNYADSVKGRFTISRDNSKNTLYLQMNSLRAEDTAVYYCARGYGSGDNWGQGTLVTVSS | EIVLTQSPGTLSLSPGERATLSCRASQSVSSSFLAWYQQKPGQAPRLLVFGASNRATGIPDRFSGSGSGTDFTLTISRLEPEDFAVYYCQQYGSSPRTFGQGTKVEIK |
| C20 | 36 | IGHV3-53*01 | IGHD6-13*01 | IGHJ4*02 | 9 | 0.3 | IGKV1-5*03 | IGKJ4*01 | 9 | 0.3 | EVQLVESGGGLIQPGGSLRLSCAASGFTVSSNYMSWVRQAPGKGLEWVSVIYSGGSTYYADSVKGRFTISRDNSKNTLYLQMNSLRAEDTAVYYCATGYSSSWRWGQGTLVTVSS | DIQMTQSPSTLSASVGDRVTITCRASQSISSWLAWYQQKPGKAPKLLIYKASSLESGVPSRFSGSGSGTEFTLTISSLQPDDFATYYCQQYNSYLLTFGGGTKVEIK |
| | 38 | IGHV3-23*01 | IGHD3-3*01 | IGHJ6*02 | 24 | 2.8 | IGLV8-61*01 | IGLJ3*02 | 10 | 0.0 | EVQLLESGGGLVQPGGSLRLSCAASGFTFSSYAMSWVRQAPGKGLEWVSGISGGGGYTYFADSVKGRFTISRDNSKNTLYLQMNSLRAEDTAIYFCAKDLAAYYEFWSGYPYFYDYGMDVWGQGTTVTVSS | QTVVTQEPSFSVSPGGTVTLTCGLSSGSVSTSYYPSWYQQTPGQAPRTLIYSTNTRSSGVPDRFSGSILGNKAALTITGAQADDESDYYCVLYMGSGIWVFGGGTKLTVL |
| | 59 | IGHV3-30-3*01 | IGHD6-19*01 | IGHJ4*02 | 15 | 1.6 | IGLV3-1*01 | IGLJ2*01 | 9 | 0.0 | EVQLVESGGGVVQPGRSLRLSCAASGFTFSSYGMHWVRQAPGKGLEWVAVISYDGSNKYYADSVKGRFTISRDNSKNTLYLQMNSLRAEDTAVYYCAKDGGFSSGWRGFDYWGQGTLVTVSS | SYELTQPPSVSVSPGQTASITCSGDKLGDKYACWYQQKPGQSPVLVIYQDSKRPSGIPERFSGSNSGNTATLTISGTQAMDEADYYCQAWDSSTVVFGGGTKLTVL |
| | 67 | IGHV1-8*01 | IGHD3-3*02 | IGHJ6*03 | 22 | 0.5 | IGKV3-15*01 | IGKJ3*01 | 8 | 0.6 | QVQLVQSGAEVKKPGASVKVSCKASGYTFTSYDINWVRQATGQGLEWMGWMNPNNGNTGYAQKFQGRVTMTRNTSISTAYMELNSLRSEDTAVYYCARGRKVPFLEWARGNYYYYMDVWGKGTTVTVSS | EIVMTQSPATLSVSPGERATLSCRASQSVSSYLAWYQQKPGQAPRLLIYGASTRATGIPARFSGSGSGTEFTLTISSLQSEDFAVYYCQQYNNWPTFGPGTKVDIR |
| | 70 | IGHV3-23*01 | IGHD2-8*01 | IGHJ4*02 | 14 | 2.5 | IGLV2-14*01 | IGLJ1*01 | 10 | 1.5 | EVQLLESGGGLAQPGGSLRLSCAASGFTFITYAMGWVRQAPGKGLEWVSSISGSGGSTYYADSVKGRFTISRDISKNTLYLQMNSLRAEDTAVYYCAKGPTIVLMVYGDYWGQGTLVTVSS | QSALTQPASVSGSPGQSITISCTGTSSDVGTYNYVSWYQQHPGKAPKLMIYDVSNRPSGVSHRFSGSKSGNTASLTISGLQAEDEADFYCSSYTSSSTLVFGTGTKVTVL |
| | 82 | IGHV4-31*03 | IGHD2-21*01 | IGHJ4*02 | 11 | 0.6 | IGKV3-15*01 | IGKJ3*01 | 9 | 0.0 | QVQLQESGPGLVKPSQTLSLTCTVSGGSISSGGYYWSWIRQHPEKGLEWIGYIYYSGNTYYNPSLKSRVTISVDTSKNQFSLKLSSVTAADTAVYYCARSDYSFLFDYWGQGTLVTVSS | EIVMTQSPATLSVSPGERATLSCRASQSVSSNLAWYQQKPGQAPRLLIYGASTRATGIPARFSGSGSGTEFTLTISSLQSEDFAVYYCQQYNNWPFTFGPGTKVDIK |
| | 119 | IGHV3-30-3*01 | IGHD2-15*01 | IGHJ5*01 | 14 | 12.9 | IGLV2-8*01 | IGLJ2*01 | 10 | 3.9 | QLHLVESGGGVVQPGRSLRLSCAASGFTFSAYNMHWVRQAPGRGLQWVAYISYRGDNKRYADPVKGRFIISRDNSENTLYLQMNNLSPEDTAVYYCAALVVVAAGDPFDSWGQGTLVTVSS | QSALTQPPSASGSPGQSVTISCTGTSSDVGGYNYVSWYQQHPGKAPKVIIYEVTKRPSGVPDRFSGSKSGNTASLTVSGLQTEDEADYYCSSYGGTNNVIFGGGTKLTVL |
| | 130 | IGHV3-30-3*01 | IGHD2-15*01 | IGHJ4*02 | 14 | 4.9 | IGLV3-10*01 | IGLJ3*02 | 11 | 0.0 | EVQLVESGGGVVQPGRSLRLSCAASGFTFSNYGMHWVRQAPGKGLEWVAVIWYDGSKKYYADSVEGRFTISRDNSRNTLYLQMNSLRAEDTALYYCAREEMVGGFNDGILWGQGTLVTVSS | SYELTQPPSVSVSPGQTARITCSGDALPKKYAYWYQQKSGQAPVLVIYEDSKRPSGIPERFSGSSSGTMATLTISGAQVEDEADYYCYSTDSSGNHRVFGGGTKLTVL |
| | 174 | IGHV3-30-3*01 | IGHD4-17*01 | IGHJ3*02 | 11 | 3.7 | IGLV1-40*01 | IGLJ3*02 | 11 | 1.6 | QVQLVESGGGVVQPGRSLRLTCAASGFTFRNYGLHWVRQAPGKGLEWVAVIWYDGSNKYYADSVKGRFTISRDNSKNTLYLQMNSLRAEDTAVYYCARKGGAYTFDIWGQGTMVTVSS | QSVLTQPPSVSGAPGQRVTISCTGSSSNIGARYDVHWYQQLPGTAPKLLLYDNSDRPSGVPDRFSGSKSGTSASLAITGLQAEDEADYYCQSYDSSLSTWVFGGGTKLTVL |
| | 192 | IGHV4-34*01 | IGHD3-10*01 | IGHJ5*02 | 18 | 1.3 | IGLV2-14*01 | IGLJ1*01 | 9 | 0.0 | QVQLQQWGAGLLKPSETLSLTCAVYGGSFSGYYWSWIRQPPGKGLEWIGEINHSGSTNYNPSLKSRVTISVDTSKNQFSLKLSSVTAADTAVYYCARGDHVSSWFGERKWFDPWGQGTLVTVSS | QSALTQPASVSGSPGQSITISCTGTSSDVGGYNYVSWYQQHPGKAPKLMIYEVSNRPSGVSNRFSGSKSGNTASLTISGLQAEDEADYYCSSYTSSLYVFGTGTKVTVL |
| | 210 | IGHV3-15*01 | IGHD2-15*01 | IGHJ2*01 | 20 | 10.0 | IGLV2-11*01 | IGLJ1*01 | 8 | 6.2 | EVQLVESGGDLVEPGGSLRLSCAASGFTFSKAWMNWVRQAPGKGLEWVGRIKSKTDGGTTDYAGPVKGRFTISRDDSKNTVYLQMDSLKTEDTAVYYCSAVRIWDCLQGICHLRNFDFWGRGNLVAVSS | QSALTQPRSVSGSPGQSVTISCTGTSSDVDGYNDVSWYQQHPGQAPKLVIYDVLKRPSGVPDRFSGFKSGSPASTAALTISDLQAADEADYYCCSYAGNYVFGPGTKVFVL |
